# Virulence strategies of an insect herbivore and oomycete plant pathogen converge on a host E3 SUMO ligase

**DOI:** 10.1101/2020.06.18.159178

**Authors:** S. Liu, C.J.G. Lenoir, T.M.M.M. Amaro, P.A. Rodriguez, E. Huitema, J.I.B. Bos

## Abstract

Pathogens and pests secrete proteins (effectors) to interfere with plant immunity through modification of host target functions and disruption of immune signalling networks. Importantly, molecular virulence strategies of distinct pathogens converge on a small set of regulators with central roles in plant immunity. The extent of convergence between pathogen and herbivorous insect virulence strategies is largely unexplored. We found that effectors from the oomycete pathogen, *Phytophthora capsici*, and the major aphid pest, *Myzus persicae* target the host immune regulator SIZ1, an E3 SUMO ligase. We show that the oomycete and aphid effector, which both contribute to virulence, feature different activities towards SIZ1. While *M. persicae* effector Mp64 increases SIZ1 protein levels, *P. capsici* effector CRN83_152 enhances SIZ1-E3 SUMO ligase activity *in vivo*. Loss of SIZ1 in host plants leads to reduced host susceptibility to aphids and an oomycete pathogen. Our results suggest convergence of distinct pathogen and pest virulence strategies on an E3 SUMO ligase to enhance host susceptibility.

## Introduction

The plant immune system is complex, featuring different classes of receptors to detect pathogens and pests and initiate multi-layered defence responses. Pattern recognition receptors (PRRs) recognize conserved pest and pathogen molecules, called pathogen associated molecular patterns (PAMPs), to activate immune responses and fight off the intruder (Jones & Dangl, 2006; Monaghan & Zipfel, 2012). Pathogens and pests deliver an arsenal of effector proteins inside their host to counter these and other plant defence pathways to promote effector-triggered susceptibility (ETS) through modulation of host protein activities. In addition, these effectors likely contribute to effective infection or infestation strategies by promoting the release of nutrients to support pathogen or pest growth. Another layer of plant immunity may be activated upon recognition of these effectors, or their activities, by nucleotide-binding leucine-rich repeat proteins (NLRs), which usually is associated with the activation of a Hypersensitive Response (HR). Given that plants carefully balance energy allocation between growth, development and reproduction, any effective immune responses need to be appropriate and controlled (Huot *et al*., 2014).

The identification of effector host targets and their effector-induced modification(s) can reveal the mechanistic basis of virulence and the biological processes that lead to susceptibility. Moreover, the identification of effector host targets for a range of pathogens pointed to convergence on key host proteins. For example, Avr2 from the fungal pathogen *Cladosporium fulvum*, EPIC1 and EPIC2B from the oomycete *Phytophthora infestans*, and Gr-VAP1 from the plant-parasitic nematode *Globodera rostochiensis* target the same defense protease Rcr3^pim^ in tomato (Song *et al*., 2009; Lozano-Torres *et al*., 2012). In addition, the effector repertoires of distinct plant pathogens, such as the bacterium *Pseudomonas syringae*, oomycete *Hyaloperonospora arabidopsidis*, and the ascomycete *Golovinomyces orontii* disrupt key components of immune signalling networks (Mukhtar *et al*., 2011; Weßling *et al*., 2014). Specifically, transcription factor TCP14 is targeted by effectors from *P. syringae*, *H. arabidopsidis* and *Phytophthora capsici*, and contributes to plant immunity (Stam *et al*., 2013c; Weßling *et al*., 2014). These findings suggest that molecular virulence strategies have evolved independently in distinct pathogens and converged on a small set of regulators with central roles in immunity.

While over the past decades our understanding of pathogen virulence strategies and susceptibility has increased dramatically, the extent with which host targets of plant-herbivorous insects overlap with other pathogens remains to be investigated. Effector biology has recently emerged as a new area in plant-herbivorous insect interactions research, leading to the identification of effector repertoires in several species (Carolan *et al*., 2009; Bos, JI *et al*., 2010; Kaloshian & Walling, 2016; Thorpe *et al*., 2018; Rao *et al*., 2019), several host targets, and insights into their contribution to the infestation process (Rodriguez *et al*., 2017; Chaudhary *et al*., 2019; Wang *et al*., 2019; Xu *et al*., 2019). These studies support an extension of the effector paradigm in plant-microbe interactions to plant-herbivorous insect interactions. Whether plant pathogenic microbes and insects adopt similar strategies to attack and reprogram their host to redirect immune responses is yet to be determined.

Here, we show that *Myzus persicae* (aphid) effector, Mp64, and *P. capsici* (oomycete) effector CRN83_152 (also called PcCRN4) (Stam *et al*., 2013a; Mafurah *et al*., 2015), associate with the immune regulator SIZ1 in the plant nucleus. SIZ1 stability and cell death activation in *N. benthamiana* are differentially affected by these effectors, suggesting these proteins feature distinct activities via this immune regulator. SIZ1 is an E3 SUMO ligase involved in abiotic and biotic stress responses, including Salicylic Acid (SA)-mediated innate immunity and EDS1/PAD4-mediated Resistance gene signalling (Miura *et al*., 2005; Catala *et al*., 2007; Lee *et al*., 2007; Miura *et al*., 2007; Jin *et al*., 2008; Miura *et al*., 2009; Ishida *et al*., 2012; Lin *et al*., 2016). Additionally, SIZ1 regulates plant immunity partially through the immune receptor SNC1 (Gou *et al*., 2017) and controls the trade-off between SNC1-dependent immunity and growth in Arabidopsis at elevated temperature (Hammoudi *et al*., 2018). By using Arabidopsis knock-out lines and gene silencing in *N. benthamiana* we show that SIZ1 negatively regulates plant immunity to aphids and an oomycete pathogen, and is required for pathogen infection and pest infestation. Moreover, Arabidopsis *siz1-2* displayed reduced susceptibility to aphids and an oomycete pathogen. Critically, the observed immunity phenotype is independent of SNC1-signalling suggesting that immunity is not specified by previously characterized SIZ1-immune signaling pathways. Our results suggest that the effector target convergence principle can be extended to herbivorous insects and raise important questions about mechanisms of action.

## Results

### Aphid effector Mp64 and oomycete effector CRN83_152 interact with AtSIZ1 and NbSIZ1

To gain novel insight into pathogen and pest effectors function towards virulence, we successfully applied yeast-two-hybrid screens to identify candidate host targets (Rodriguez *et al*., 2017). We identified the E3 SUMO ligase SIZ1 in screens against a *N. benthamiana* library (generated from aphid infested and *P. capsici* infected leaves) with *M. persicae* (aphid) effector Mp64 and *P. capsici* (oomycete) effector CRN83_152 as baits. Mp64 was screened against an estimated 5×10^6^ cDNAs and revealed 2 independent prey clones with an insert showing similarity to SIZ1, whilst the effector CRN83_152 screen of 4×10^6^ yeast transformants, identified 3 independent prey clones with an insert similar to SIZ1. All putative interactors identified in the two effector screens are summarized in Table S1. Since all NbSIZ1 (*N. benthamiana* SIZ1) prey clones from the Mp64 and CRN83_152 screens were partial-length, we designed primers to amplify and clone the full-length NbSIZ1. Although we were unable to amplify NbSIZ1 based on the two best BLAST hits against the *N. benthamiana* genome (Niben101Scf15836g01010.1 and Niben101Scf04549g09015.1) due to poor/no primer annealing at the 3’ end, we successfully amplified NbSIZ1 sequences based on the 3’ end of SIZ1 sequences from *N. attenuata* (XP_019237903) and *N. tomentosiformis* (XP_018631066). The full-length NbSIZ1 sequence we cloned was identical to our partial yeast prey clones and NbSIZ1 database sequences, except for a 27 amino acid insertion at position 225-252. A direct comparison between NbSIZ1 and the well characterised AtSIZ1 showed 60% identity between proteins (Fig.S1). Given that AtSIZ1 is well characterised and helps regulate plant immunity (Miura *et al*., 2005; Catala *et al*., 2007; Lee *et al*., 2007; Hammoudi *et al*., 2018), we included AtSIZ1 in our efforts to further validate effector-SIZ1 interactions and characterise the role of SIZ1 in plant-aphid/oomycete interactions using both Arabidopsis and *N. benthamiana* resources. It should be noted that whilst Arabidopsis is a host for *M. persicae*, this plant species is not a natural host for *P. capsici*. We first tested whether Mp64 and CRN83_152 interact with full-length NbSIZ1 and AtSIZ1 in yeast. Whilst yeast reporter assays showed interaction of Mp64 with both full-length SIZ1 versions (Fig. S2), we were unable to obtain yeast co-transformants expressing both CRN83_152 and full-length SIZ1 in repeated transformation experiments that included transformation controls. We also included a mutant of CRN83_152, called CRN83_152_6D10 (Amaro *et al*., 2018), which does not trigger CRN-cell death activation but retains virulence activity, in co-transformation experiments with similar results. Based on further data presented below, we hypothesize that the lack of CNR83_152/SIZ1 yeast co-transformants is due to enhanced E3 SUMO ligase activity of SIZ1 in the presence of this effector which may affect yeast cell viability.

To test for *in planta* effector-SIZ1 interactions, we co-expresssed GFP-Mp64 and GFP-CRN83_152_6D10 with either AtSIZ1-myc or NbSIZ1-myc in *N. benthamiana* (Fig. 1a,b). Immunoprecipitation of both effectors resulted in the co-purification of NbSIZ1, suggestive of an association *in planta* (Fig. 1b). Co-immunoprecipitation of the effectors with AtSIZ1 gave similar results, however, with CRN83_152 and CRN83_152_6D10 only showing a weak band corresponding to AtSIZ1 upon co-purification (Fig. 1a). Altogether our data demonstrate that effectors from two distinct plant parasites associate with the same host protein, SIZ1, *in planta*, prompting us to further investigate the contribution of SIZ1 and the effectors to susceptibility.

**Fig. 1.**
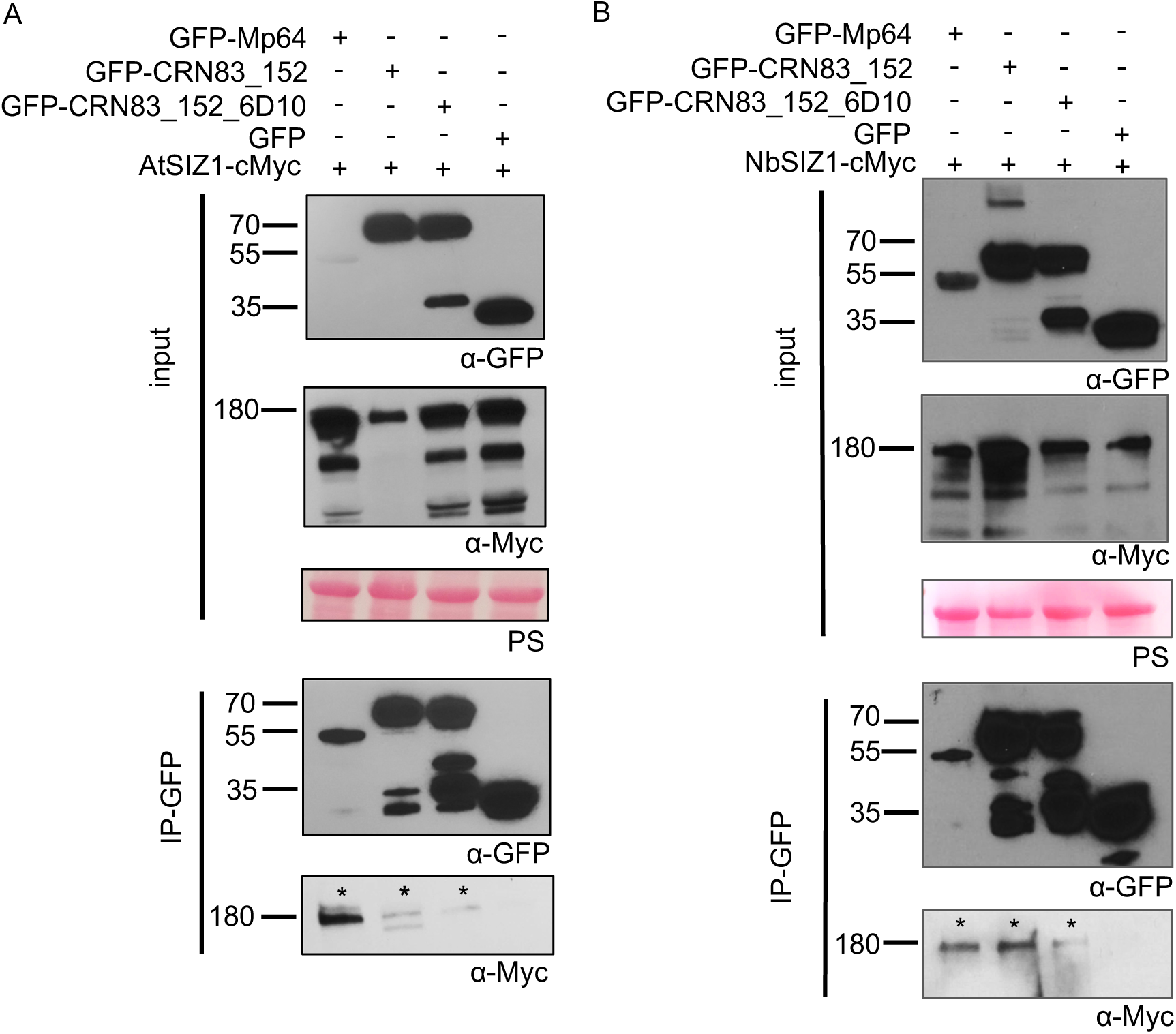
*Myzus persicae* effector Mp64 and *Phytophthora capsici* effector CRN83_152 associate with SIZ1 from Arabidopsis and *Nicotiana benthamiana*. (A) Immunoprecipitation (IP) of protein extracts from agroinfiltrated leaves using GFP-Trap confirmed that AtSIZ1-cMyc(10x) associates with GFP-Mp64, GFP-CRN83_152 and GFP-CRN83_152_6D10, but not with the GFP control. (B) Immunoprecipitation (IP) of protein extracts from agroinfiltrated leaves using GFP-Trap confirmed that NbSIZ1-cMyc(10x) associates with GFP-Mp64, GFP-CRN83_152 and GFP-CRN83_152_6D10, but not with the GFP control. Protein size markers are indicated in kD, and protein loading is shown upon ponceau staining of membranes. Experiments were repeated at least three times with similar results.

### Silencing of *NbSIZ1* reduces *N. benthamiana* host susceptibility to *P. capsici*

To assess the contribution of SIZ1 to immunity in a *P. capsici* host species, we made use of Virus Induced Gene Silencing (VIGS) in *N. benthamiana* (Ratcliff *et al*., 2001; Lu *et al*., 2003). Our TRV-*NbSIZ1* construct, designed to silence *NbSIZ1*, reduced transcripts levels by around 60% compared with plants expressing the TRV-*GFPfrag* control (a fragment of GFP) (Fig S3). Silenced plants showed a slight reduction in growth compared with the TRV-*GFPfrag* control and cell death in older leaves (Fig. S3). In our hands, VIGS assays based on TRV in *N. benthamiana* are incompatible with aphid assays (TRV infection causes aphids to die), therefore, we only performed infection assays with *P. capsici* on *NbSIZ1* silenced plants. Detached leaves were used for *P. capsici* infection assays based on zoospore droplet inoculations, followed by lesion size diameter measurements. *P. capsici* lesion size on *NbSIZ1* silenced leaves was significantly reduced 2-4 days after inoculation when compared to control plants (Mann-Whitney U test, p<0.0001; Fig. 2; Fig. S3). These results indicate that NbSIZ1 contributes to host susceptibility to this oomycete plant pathogen.

**Fig. 2.**
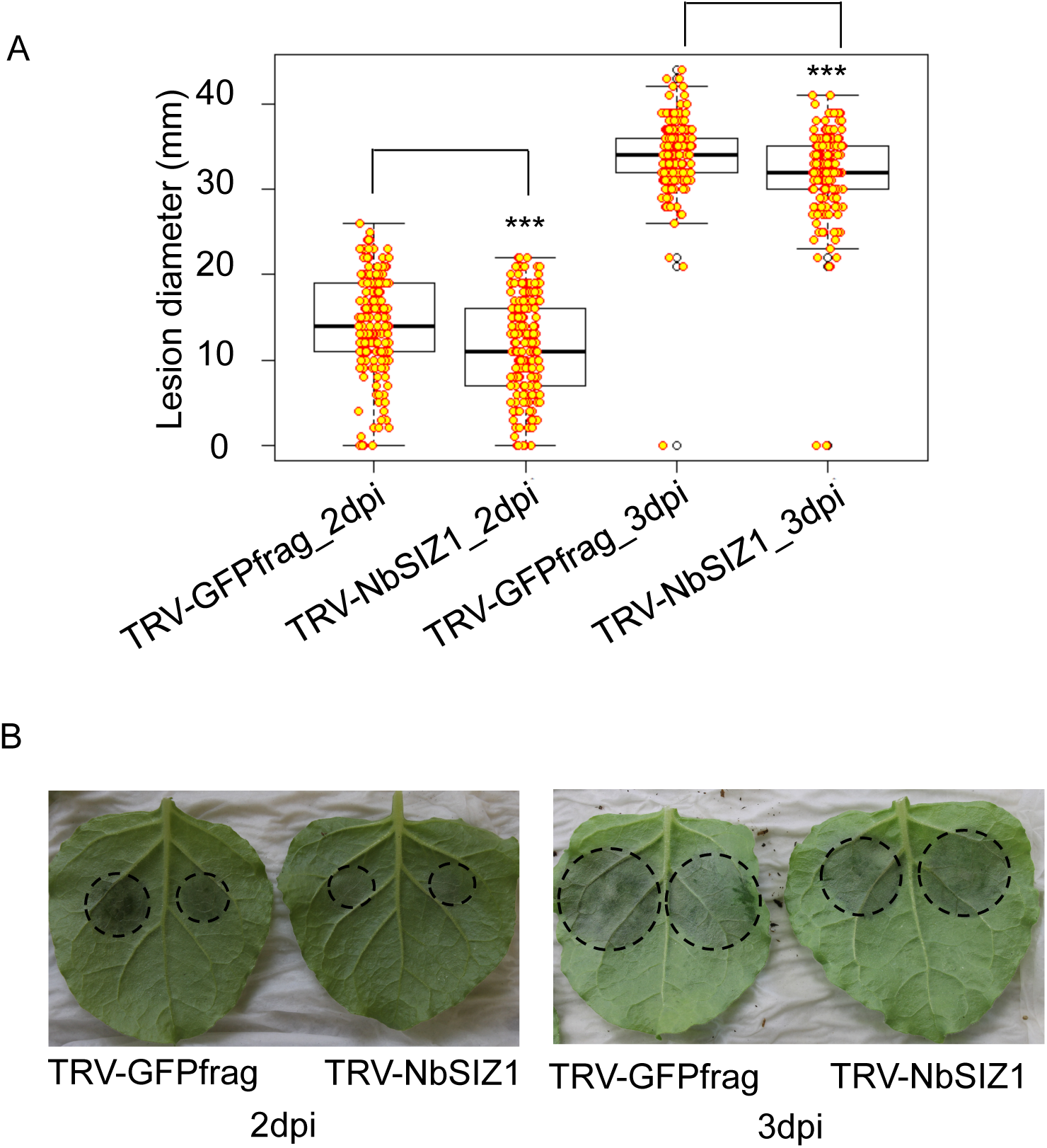
Virus-induced gene silencing of *NbSIZ1* reduces host susceptibility to *Phytophthora capsici*. (A) Boxplot showing the lesion diameter of *P. capsici* infection sites on *Nicotiana benthamiana* control (TRV-*GFPfrag*) or *NbSIZ1*-silenced plants (TRV-*NbSIZ1*). 5ul of zoospore suspension (50,000 spores/ml) was drop-inoculated on *N. benthamiana* leaves. Data was collected 2 and 3 days post inoculation (dpi) from three biological replicates (n=48 per biological replicate). Asterisks denote significant difference between the *GFPfrag* control and *NbSIZ1*-silenced plants (Mann-Whitney U test, p<0.0001). (B) Representative images of *NbSIZ1*-silenced and *GFPfrag* control leaves 2 days and 3 days post inoculation (dpi) with *P. capsici* zoospores.

### Loss-of-function mutation *siz1-2* in Arabidopsis leads to reduced susceptibility to *M. persicae* and *P. capsici*

Since AtSIZ1 negatively regulates plant innate immunity in Arabidopsis to the bacterial plant pathogen *Pseudomonas syringae* pv. Tomato DC3000 (Pst) (Lee *et al*., 2007), we tested whether this also applies to interactions with *M. persicae* and *P. capsici*. We performed aphid performance assays, based on fecundity measurements, as well as *P. capsici* infection assays on the Arabidopsis loss!of-function mutant *siz1-2*. Given that siz1-2 mutants have a dwarf phenotype, associated with SA hyper-accumulation, we included Arabidopsis line *siz1-2*/NahG, in which this phenotype is (partially) abolished (Lee *et al*., 2007). While Arabidopsis is a host for the aphid *M. persicae,* only few *P. capsici* isolates infect Arabidopsis under controlled environmental conditions and high levels of inoculum (Wang *et al*., 2013), suggesting that Arabidopsis is not a natural host. Aphid performance assays showed a significant reduction in fecundity on the *siz1-2* and *siz1-2*/*NahG* lines compared to the Col-0 (ANOVA, p<0.0001; Fig. 3a) and *NahG* controls (ANOVA, p<0.01; Fig. 3a), respectively, with only few aphids surviving on the *siz1-2* line. The *siz1-2* reduced susceptibility to aphids is largely maintained in the *NahG* background, implying that this phenotype is largely independent of SA accumulation.

**Fig. 3.**
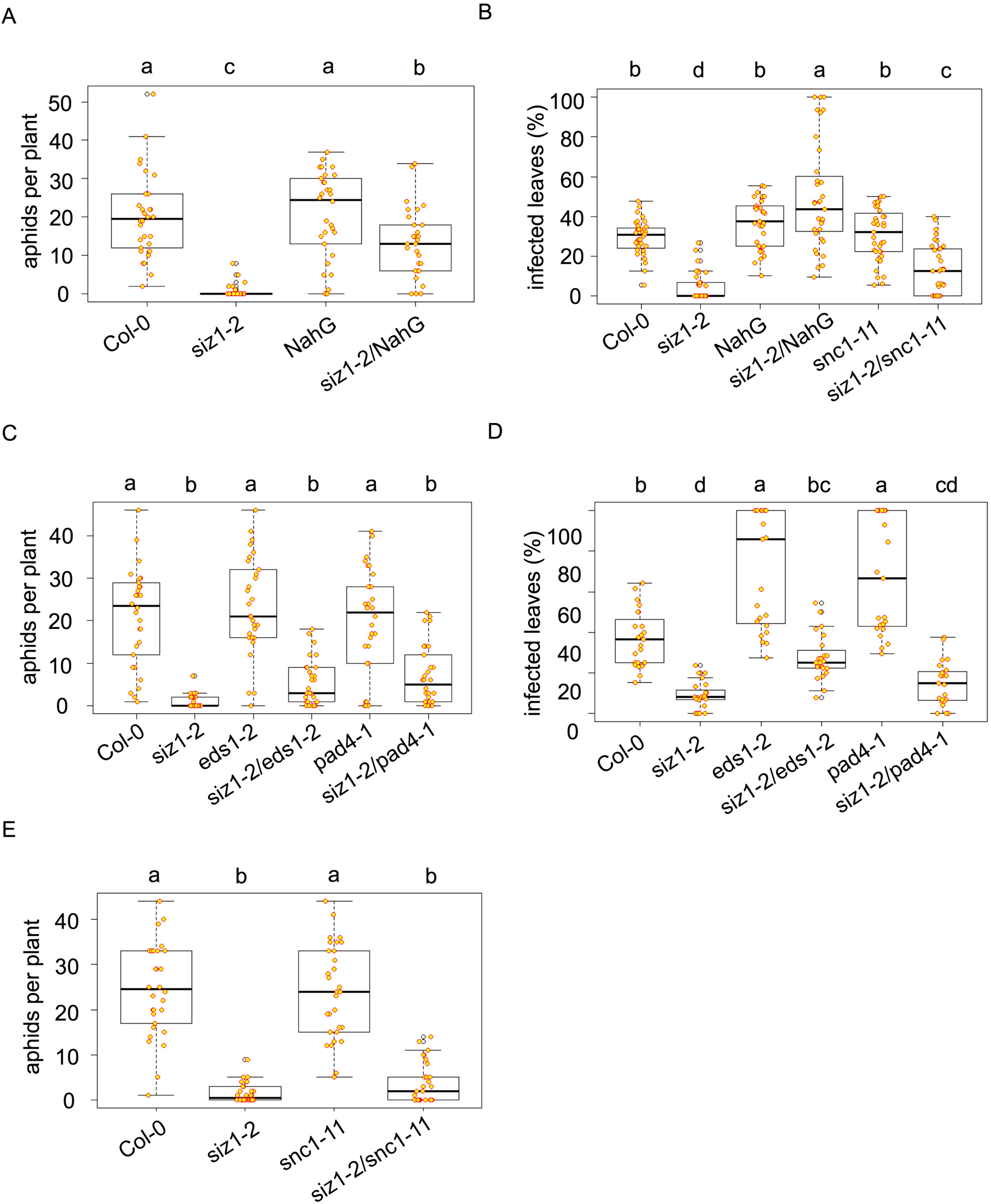
The Arabidopsis *siz1-2* mutant shows reduced susceptibility to *Myzus persicae* and enhanced resistance to *Phytophthora capsici*. For aphid infestation assays (A, C, E) plants were infested with two 2-day old nymphs and the aphid populations were counted 10 days later. For *P. capsici* infection assays (B, D) plants were spray-inoculated with a zoospore suspension of 100,000 spores/ml and the percentage of symptomatic leaves was recorded 8 days later. A linear mixed effects model with experimental block and biological replicate as random factors was fitted in dataset from aphid fecundity assays. A linear mixed model with biological replicate as random effect was used in dataset of *P. capsici* infection assays. ANOVA was used to analyse the final models and a post-hoc test was performed by calculating the Least Squares Means by using an emmeans package in R. Different letters denote significant differences within a set of different plant genotypes. (A) Arabidopsis mutant *siz1-2* is less susceptible to *M. persicae* (aphids) than the Col-0 control with significantly less aphids recorded on the mutant versus control plants (p<0.0001), including in the *NahG* background (p<0.01). (B) Arabidopsis mutant *siz1-2* shows enhanced resistance to *P. capsici* compared with the Col-0 control, with significantly less infected leaves on the mutant versus control plants (p<0.0001). In the *NahG* background, *siz1-2* is associated with increased infection compared to the *NahG* control (p<0.01). The *siz1-2/snc1-11* mutant shows enhanced resistance to *P. capsici* compared with the *snc1-11* mutant, with significantly less leaves infected on the double mutant (p<0.0001). (C) Arabidopsis mutants *siz1-2/esd1-2* and *siz1-2/pad4-1* are less susceptible to *M. persicae* (aphids) than the *eds1-2* and *pad4-1* mutants, respectively, with significantly less aphids recorded on the double compared with the single knock-out mutants (p<0.0001). (D) Arabidopsis mutants *siz1-2/eds1-2* and *siz1-2/pad4-1* are more resistant to *P. capsici* than the *eds1-*2 and *pad4-1* mutants, respectively, with significantly less leaves infected on the double compared with the single knock-out mutants (p<0.0001). (E) Arabidopsis mutant *siz1-2/snc1-11* is less susceptible to *M. persicae* (aphids) than the compared *snc1-11* mutant, with significantly less aphids recorded on the double mutant (p<0.0001).

For *P. capsici* infection assays, plants were spray inoculated with a zoospore solution and the percentage of symptomatic leaves was counted 10 days later. The percentage of symptomatic *siz1-2* leaves was reduced by 83% compared with the Col-0 control (ANOVA, p<0.0001; Fig. 3b, Fig. S4). We did not observe a difference in *P. capsici* infection levels between the *NahG* line and Col-0 but did note a slight increase in infection on *siz1/NahG* compared to the *NahG* background (ANOVA, p<0.01; Fig. 3b, Fig. S4).

### Arabidopsis s*iz1-2* reduced susceptibility to *M. persicae* and *P. capsici* is independent of SNC1, EDS1 and PAD4

EDS1, PAD4 and SNC1 are required for *siz1-2* enhanced resistance to *P. syringae* pv tomato DC3000 (Lee *et al*., 2007; Gou *et al*., 2017; Hammoudi *et al*., 2018). To explore whether these signalling components also contribute to reduced aphid infestation and *P. capsici* infection on *siz1-2,* we performed aphid infestation and infection assays on Arabidopsis *siz1-2/eds1-2*, *siz1-2/pad4-1* and *siz1-2/snc1-11* double mutants.

Aphid infestation on the *siz1-2/eds1-2* mutant was reduced by 75% compared with the *eds1-2* mutant (ANOVA, p<0.0001, Fig. 3c), and was comparable to *siz1-2* (Fig. 3c), suggesting that the reduced susceptibility of *siz1-2* to aphids is independent of EDS1. In addition, aphid fecundity was reduced on *siz1-2/pad4-1* by around 65% compared with the *pad4-1* mutant (ANOVA, p<0.0001, Fig. 3c), and was comparable to *siz1-2*. These data suggest that *siz1-2* reduced susceptibility to aphids is also independent of PAD4.

In line with previous reports (Wang *et al*., 2013) the *eds1-2* and *pad4-1* mutants were less resistant to *P. capsici* than Col-0 (ANOVA, p<0.0001, Fig. 3d, Fig. S4), indicating EDS1 and PAD4 contribute to Arabidopsis nonhost resistance to this pathogen. The percentage of symptomatic *siz1-2/eds1-2* and *siz1-2/pad4-1* leaves was around 60% and 55% less compared to the *eds1-2* (ANOVA, p<0.0001, Fig. 3d, Fig. S4) and *pad4-1* (ANOVA, p<0.0001, Fig. 3d, Fig. S4) mutants, respectively. Similar to our aphid data, Arabidopsis *siz1-2* enhanced resistance to *P. capsici* was maintained in the *eds1-2* and *pad4-1* mutant backgrounds and when compared to the appropriate controls (*eds1-2* and *pad4-1*, respectively).

Aphid fecundity on *siz1-2/snc1-11* was approximately 85% reduced compared with the *snc1-11* control (ANOVA, p<0.0001), and was comparable to *siz1-2* (Fig. 3e), suggesting that *siz1-2* reduced susceptibility to aphids is independent of SNC1. The *siz1-2/snc1-11* double mutant also showed enhanced resistance to *P. capsici*, with 55% less symptomatic leaves compared to the *snc1-11* mutant (ANOVA, p<0.0001, Fig. 3b, Fig. S4). The percentage of symptomatic leaves on *siz1-2/snc1-11* was slightly higher compared with the *siz1-2* mutant (ANOVA, p<0.05, Fig. 3b, Fig. S4). With the *siz1-2* enhanced resistance to *P. capsici* largely maintained in the *snc1-11* background, this phenotype is likely independent of the immune receptor SNC1. Overall, *siz1-2* reduced susceptibility to both *M. persicae* and *P. capsici* is independent of defence signalling components previously implicated in SIZ1 immune functions. These data are in line with a model wherein SIZ1 contributes to host susceptibility to certain pests and pathogens, perhaps upon effector-mediated modulation.

### Nuclear aphid effector Mp64 enhances Arabidopsis susceptibility to *M. persicae*

While the nuclear PcCRN83_152 effector from *P. capsici* was previously shown to be essential for pathogen virulence and promotes plant susceptibility (Stam *et al*., 2013a; Mafurah *et al*., 2015), the role of aphid effector Mp64, which was identified as a candidate effector in *A. pisum* and *M. persicae* through bioinformatics pipelines (Carolan *et al*., 2011; Thorpe *et al*., 2016; Boulain *et al*., 2018), is unknown. Mp64 is a protein of unknown function with a predicted nuclear localisation based on ProteinPredict (Yachdav *et al*., 2014) and NLStradamus (Nguyen Ba *et al*., 2009), and Mp64 homologs are present in other aphid species (Fig. S5). We investigated the subcellular localisation of Mp64 by confocal microscopy of *N. benthamiana* leaves transiently expressing GFP-Mp64 (lacking the predicted signal peptide). Imaging of epidermal cells expressing Mp64 revealed accumulation of GFP-Mp64 in the nucleus and nucleolus, with no signal detectable in the cytoplasm (Fig. 4a). Nuclear localisation of GFP-Mp64 was confirmed upon co-localisation with the nuclear marker Histone 2B (H2B) (Fig. 4b). In addition, we observed dots within the nucleoplasm corresponding to GFP-Mp64.

**Fig. 4.**
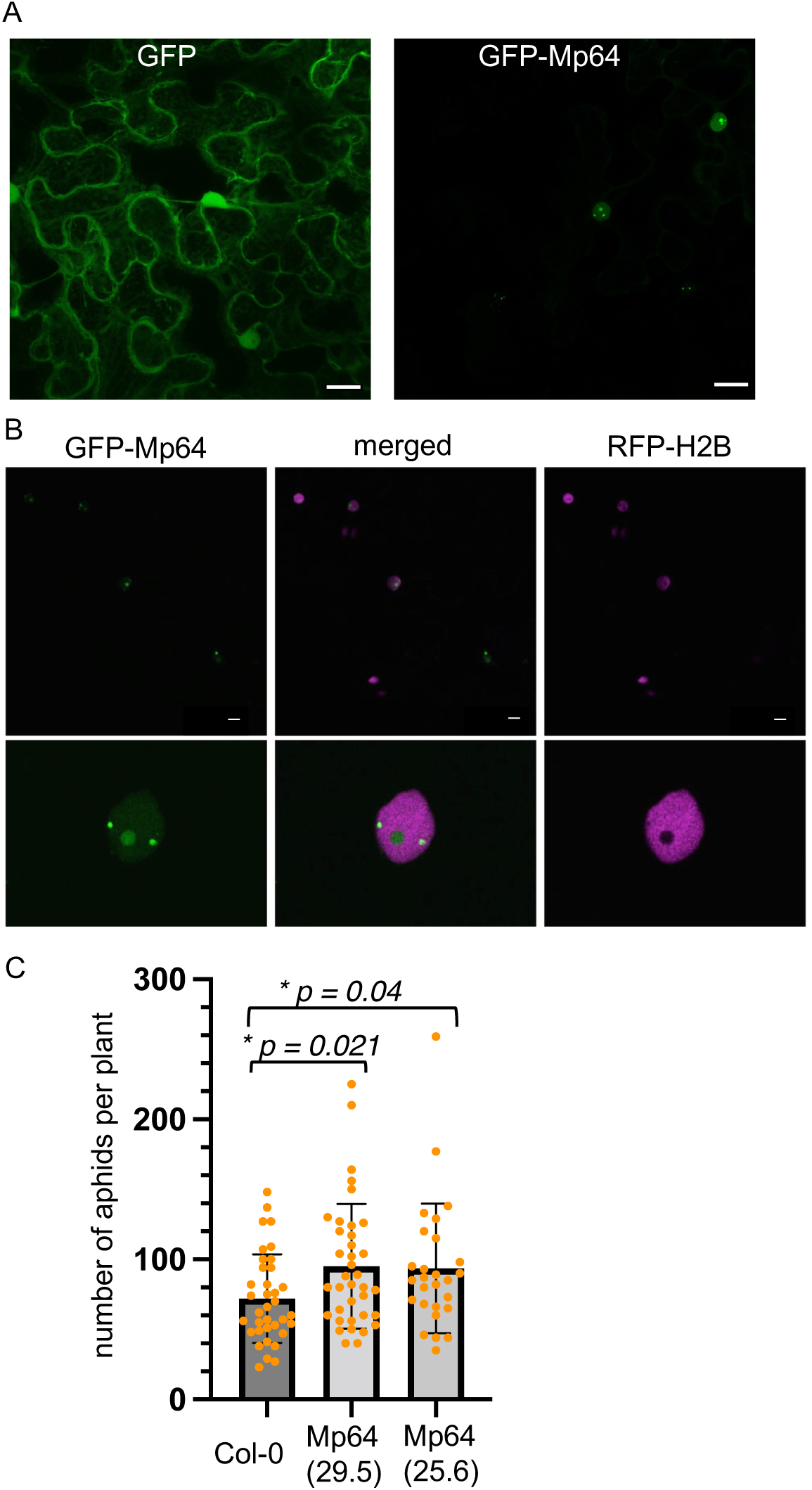
Constitutive ectopic expression of nuclear aphid effector Mp64 in Arabidopsis enhances susceptibility to *Myzus persicae*. (A) Nuclear localisation of aphid effector Mp64 in *Nicotiana benthamiana* with free GFPas a control. (B) Confocal images showing nuclear localisation of aphid effector Mp64 in host Nicotiana benthamiana. Leaves transiently expressing GFP-Mp64 and histone marker RFP-Histone 2B (H2B), a nuclear marker, were used for confocal imaging 2 days after agroinfiltration. Images in the upper panel were collected as z-stack projection and those in the lower panel were collected as single optical sections through nuclei of cells ectopically overexpressing Mp64. Scale bar represents 10µm. (C) Two Arabidopsis transgenic lines, 25.6 and 29.5, were challenged with two apterous adult aphids. Total numbers of aphids per plant were counted 10 days post infestation. The boxplot displays the distribution of datapoints from three independent biological replicates (n=10 per replicate). The black line within the box represents the median. The top and bottom edges of the box indicate upper quantile and lower quantile. The datapoints out of the upper and lower extreme of the whisker are outliners. Asterisks denote significant difference between treatments and control, with corrected p-values indicated (Kruskal Wallis, multiple comparison Benjamini-Hochberg correction based on FDR).

To confirm that Mp64 contributes to aphid virulence, we generated Arabidopsis transgenic lines expressing the mature Mp64 protein driven by the 35S promoter. Arabidopsis lines expressing Mp64 showed no developmental or growth phenotypes (Fig. S6) and were subjected to aphid fecundity assays. Two age-synchronized *M. persicae* aphids were placed on transgenic Mp64 lines and Col-0 control plants, and progeny was counted after 10 days. The average number of aphids on two independent transgenic Mp64 lines (25.6 and 29.5) was around 30% higher than on the Col-0 control (Kruskal-Wallis test with multiple comparisons Benjamini-Hochberg correction based on FDR) Fig. 4c), indicating that Mp64 enhances Arabidopsis host susceptibility to *M. persicae*. While our two Mp64 transgenic lines showed different levels of Mp64 expression, with line 29.5 showing lower expression than line 25.6. We did not find a correlation between higher expression and virulence effect, with both lines showing a similar increase in aphid numbers.

To test whether Mp64 also affects *P. capsici* infection, we transiently over-expressed Mp64 and a vector control in *N. benthamiana* and challenged infiltration sites with a zoospore suspension. While *P. capsici* effector CRN83_152 enhanced *N. benthamiana* susceptibility to *P. capsici*, in line with previous reports (Stam *et al*., 2013b) (Mafurah *et al*., 2015; Amaro *et al*., 2018). Aphid effector Mp64 did not affect host susceptibility to *P. capsici* (Fig. S7), pointing to distinct virulence activities of these effectors.

### Distinct nuclear localization patterns of Mp64, CRN83_152 and SIZ1

Since both Mp64 and CRN83_152 are nuclear effectors, and their host interacting protein SIZ1 is reported to localise and function in the plant nucleus (Miura *et al*., 2005), we determined whether the effectors co-localise with SIZ1 in this subcellular compartment. We performed confocal imaging of *N. benthamiana* leaves transiently co-expressing SIZ1-mRFP and GFP-effector fusions. We did not detect signal corresponding to the effectors or SIZ1 outside the nucleus (Fig. 4; Fig. 5; Fig S8). In line with previous reports, mRFP signal corresponding to both AtSIZ1 and NbSIZ1 was visible in the plant nucleoplasm, along with distinct speckles in the nucleolus (Fig. 5; Fig. S8). Expression of full-length SIZ1-RFP was confirmed by Western blotting (Fig. S9). GFP signal corresponding to GFP-Mp64 and CRN83_152 was detectable in the nucleus, with GFP-Mp64 more localized within the nucleolus and in speckles within the nucleoplasm, and GFP-CRN83_152 present only in the nucleoplasm (Fig. 4; Fig. 5; Fig S8). Indeed, GFP-CRN83_152 co-localises with AtSIZ1-RFP and NbSIZ1-RFP in the nucleoplasm, whereas GPF-Mp64 shows a more distinct localization in the nucleus that only partially overlaps with nucleoplasm localization of SIZ1. Interestingly, nucleolar speckles corresponding to SIZ1-RFP occasionally coincided with loss of GFP-Mp64 signal, and GFP-Mp64 speckles within the nucleoplasm coincided with loss of SIZ1-RFP signal. However, we did not find evidence for altered localization of SIZ1 in the presence of the effectors, and vice versa (Fig. 5; Fig. S9). With the effectors and SIZ1 only detected within the nucleus, this likely is the compartment where interactions take place.

**Fig. 5.**
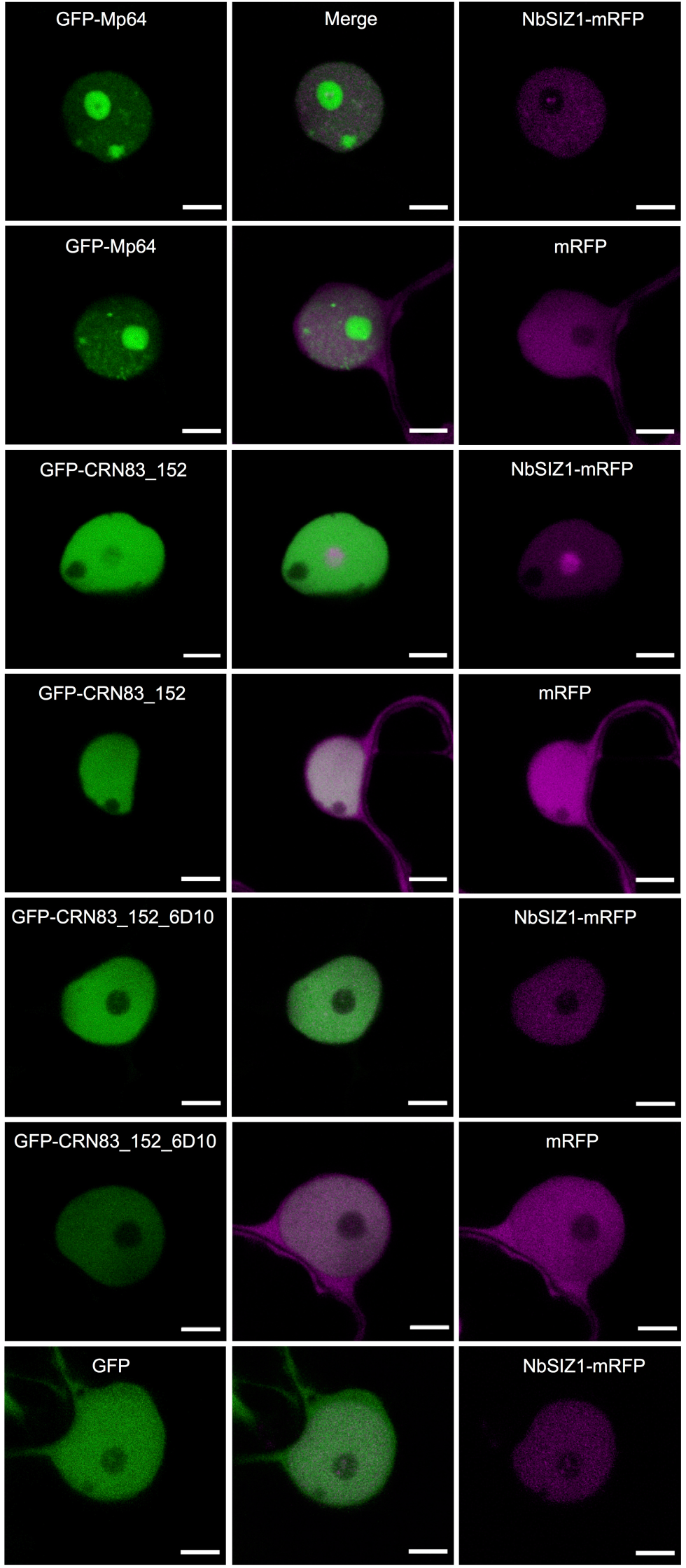
Localisation of effectors Mp64, CRN83_152(6D10 and NbSIZ1 in the host nucleus. Leaves transiently expressing GFP-Mp64, GFP-CRN83_152 or GFP-CRN83_152_6D10 in combination with RFP or NbSIZ1-RFP were used for confocal imaging around 36 hours after agroinfiltration. Images show single optical sections through nuclei co-expressing the GFP-effector with NbSIZ1-RFP or RFP control. Scale bars represent 5 µm.

### Mp64 but not CRN83_152_6D10 stabilises AtSIZ1 in planta

In co-immunoprecipitation experiments, we consistently observed increased protein levels of AtSIZ1 in input samples upon co-expression with Mp64 but not CRN83_152_6D10 (Fig 1a,b). To test whether Mp64 indeed stabilizes SIZ1 *in planta*, we performed co-expression assays of both effectors with SIZ1 in parallel in three independent experiments. Western blot analyses combined with quantitative analyses of band intensities, showed that AtSIZ1 levels were higher than NbSIZ1 levels among all replicates. In addition, we consistently observed an increase in SIZ1 protein levels in the presence of Mp64 compared to the GFP-GUS control (Fig. 6a, additional replicates in Fig. S10), indicating that Mp64 stabilizes SIZ1 *in planta*. Although not consistent across replicates, we did observe slightly decreased AtSIZ1 levels in the presence of CRN83_152_6D10, compared to the GFP control, but this observation was not consistent across replicated experiments (Fig 6a, additional replicates in Fig S10).

**Fig. 6.**
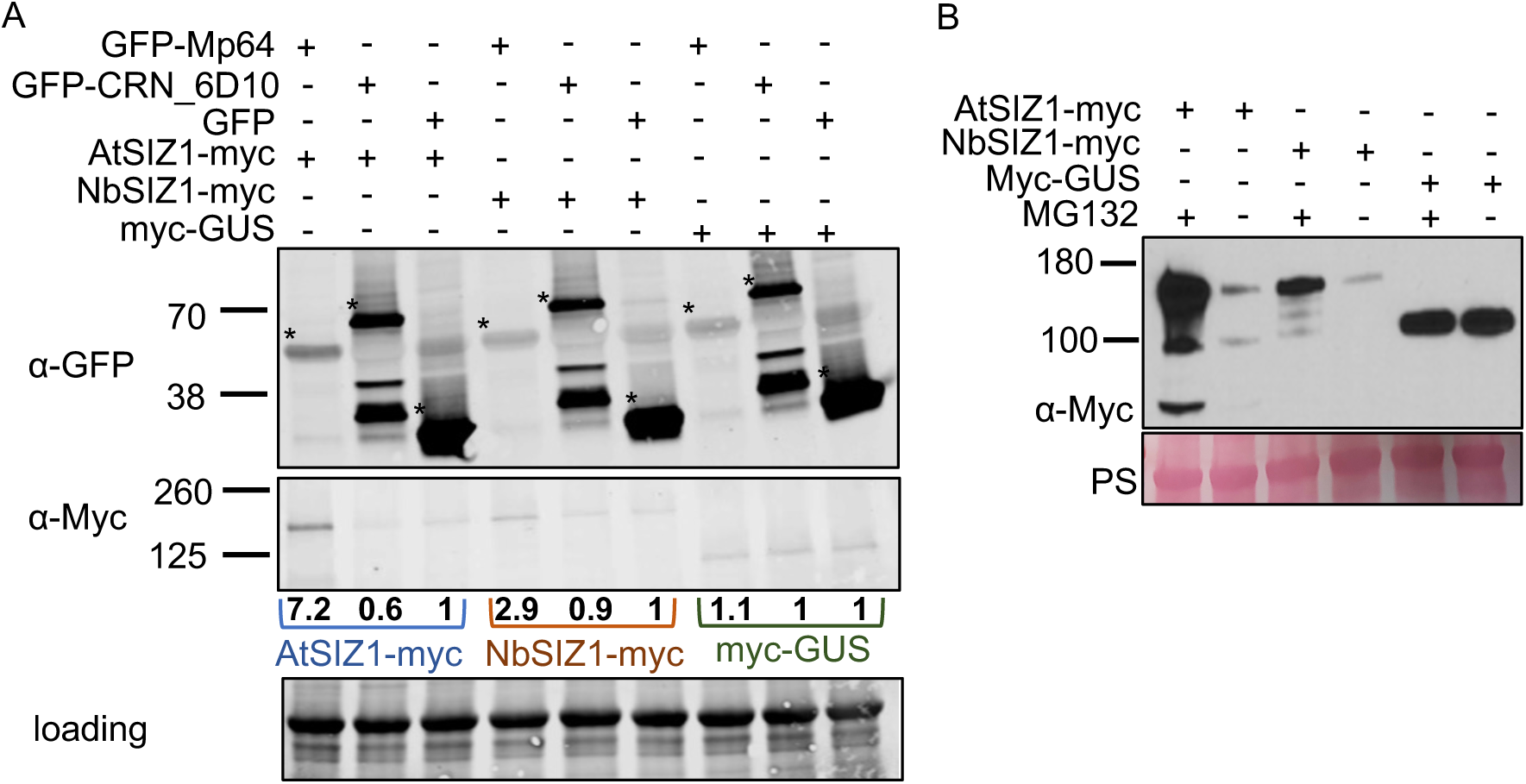
Mp64 but not CRN83-152_6D10 enhanced SIZ1 stability *in planta*. (A) Western blots showing Mp64 increases SIZ1 protein levels. Blots were prepared using total plant extracts of *N. benthamiana* infiltration sites expression GFP-Mp64/CRN83-152_D610 or GFP (control) with SIZ1-myc. Leaf material was harvested 2 dpi. Total protein amount was detected using the Revert™ 700 Total Protein Stain followed by imaging in the 700 nm channel using an Odyssey^®^ CLx Imaging System. The panel indicated by “loading” shows a proportion of the membrane that includes Rubisco. Detection of GFP-fusion and SIZ-myc fusion proteins upon antibody incubation was in the 800nm channel using an Odyssey^®^ CLx Imaging System. Asterisk indicated bands corresponding to GFP-effectors/GFP. SIZ1 protein quantitation was done by normalizing the band intensity of SIZ1-myc to the total protein amounts using Empiria Studio 2.1 (Licor). SIZ1-myc levels in samples with GFP-Mp64 or GFP-CRN83_152_6D10 were compared to GFP (control, set at 1) to generate band intensity ratios, indicated by values below the western blot incubated with Myc-antibodies. (B) Western blot showing the 26S proteasome inhibitor MG132 increases detectable levels of SIZ1. Protein extracts were prepared from *N. benthamiana* leaves expressing SIZ1-myc or GUS, challenged with MG132 treatment, and used for western blotting. Ponceau staining was used to show protein loading (PS), and myc-antibodies were used to detect SIZ1-myc across samples.

Overall, we noted that full length AtSIZ1 and NbSIZ1 are rather difficult to detect by western blotting, suggesting low expression and/or low protein stability. Indeed, Lin et al (Lin *et al*., 2016) previously showed that the 26S proteasome inhibitor MG132 reduces degradation of GFP-AtSIZ1 mediated by the ubiquitin E3 ligase COP1 (CONSTITUTIVE PHOTOMORPHOGENIC 1), an ubiquitin E3 ligase. In line with this we show that both AtSIZ1 and NbSIZ1 are more readily detected by Western blotting upon MG132 treatment (Fig 6b), suggesting that the levels of both these SIZ1 versions is tightly regulated *in planta*.

### CRN83_152_6D10 but not Mp64 enhances AtSIZ1 triggered cell death in *N. benthamiana*

When performing transient expression assays in *N. benthamiana* with AtSIZ1 and NbSIZ1, we observed the onset of cell death starting from 3 days after infiltration specifically upon expression of AtSIZ1. We investigated whether co-expression of the aphid and oomycete effectors with SIZ1 would either enhance or reduce this cell death activation. In the absence of any effectors, AtSIZ1 consistently activated cell death from 3-4 days after infiltration, whereas only occasional microscopic cell death was visible in infiltration sites expressing NbSIZ1. Both AtSIZ1 and NbSIZ1 fusion proteins, with a C-terminal RFP tag, were detectable in transient expression assays (Fig. S9). While co-expression of Mp64 with SIZ1 did not affect the cell death phenotype, co-expression of CRN83_152_6D10 with AtSIZ1 led to a stronger cell death response compared to the AtSIZ1 and CRN83_152_6D10 controls (Fig. 7; Fig. S11). These data suggest that CRN83_152_6D10 but not Mp64 enhances AtSIZ1-triggered cell death.

**Fig. 7.**
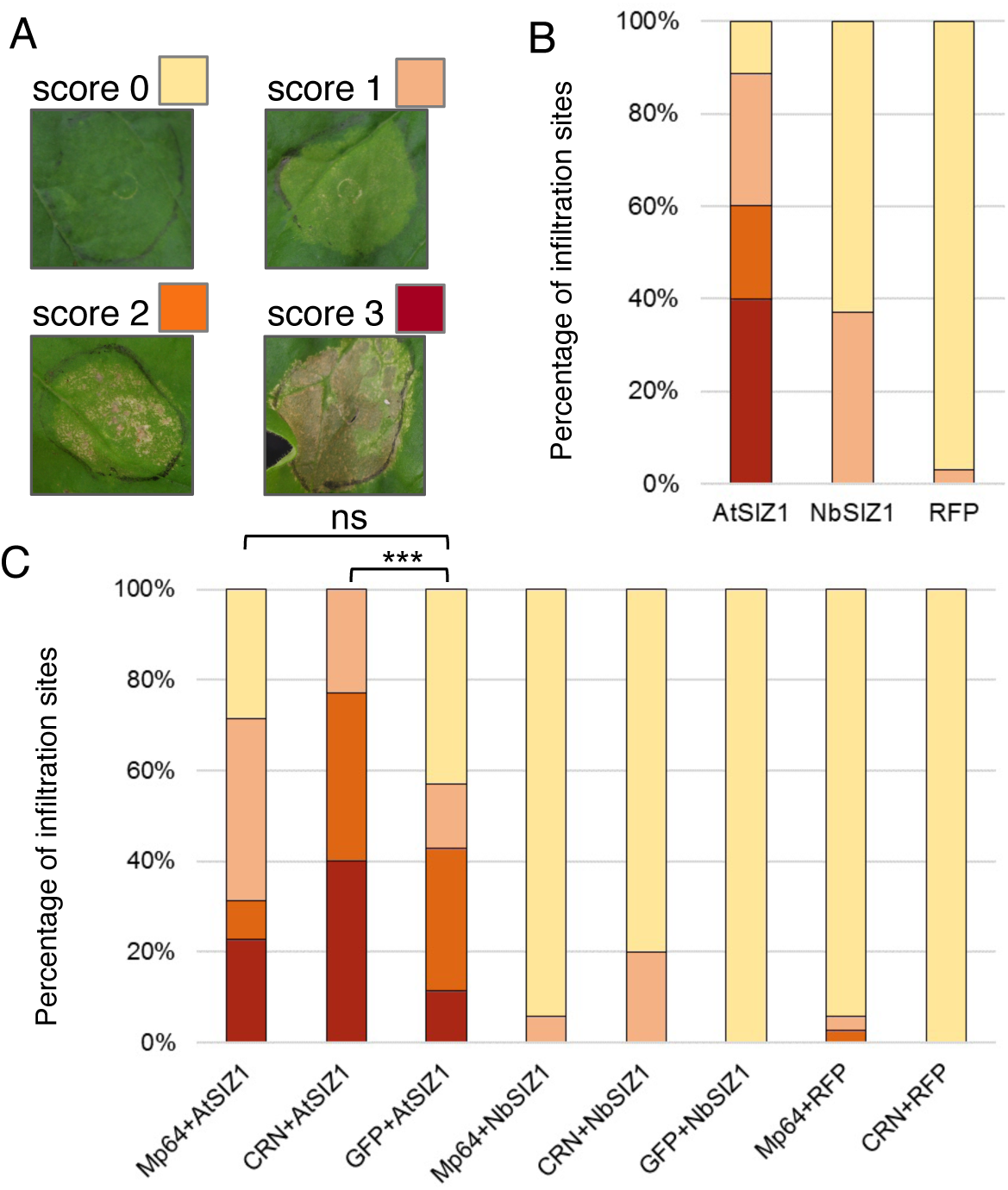
SIZ1-triggered cell death in *N. benthamiana* is enhanced by CRN83_152_6D10 but not Mp64. (A) Scoring overview of infiltration sites for SIZ1-triggered cell death. Infiltration site were scored for no symptoms (score 0), chlorosis with localized cell death (score 1), less than 50% of the site showing visible cell death (score 2), more than 50% of the infiltration site showing cell death (score 3). (B) Bar graph showing the proportions of infiltration sites with different levels of cell death upon expression of AtSIZ1, NbSIZ1 (both with a C-terminal RFP tag) and an RFP control. Graph represents data from a combination of 3 biological replicates of 11-12 infiltration sites per experiment (n=35). Data was collected 7 days after infiltration. (C) Bar graph showing the proportions of infiltration sites with different levels of cell death upon expression of SIZ1 (with C-terminal RFP tag) either alone or in combination with aphid effector Mp64 or Phytophthora capsica effector CRN83_152_6D10 (both effectors with GFP tag), or a GFP control. Data was collected 7 days after infiltration. Graph represent data from a combination of 3 biological replicates of 11-12 infiltration sites per experiment (n=35). *** indicates p < 0.001 (Kruskal-Wallis test with post-hoc Dunn’s test for multiple comparison,) and ns indicates no significant difference.

### CRN83_152 but not Mp64 enhances SUMOylation upon transient co-expression with SIZ1

To assess whether both AtSIZ1 and NbSIZ1 are active upon transient expression in *N. benthamiana* and whether effectors CRN83_152 and Mp64 alter any E3 SUMO ligase activity, we performed co-expression assays with RFP-tagged AtSUMO1. First, we co-infiltrated *Agrobacterium* strains carrying constructs for AtSIZ1-myc, NbSIZ1-myc and myc-GUS (control) with RFP-SUMO1, to assess whether ectopic/overexpression of SIZ1 increased detectable SUMO profiles upon heat treatment. Western blot analyses showed an increase in the presence of SUMO profiles, as detected with RFP-antibodies against RFP-SUMO1, upon expression of SIZ1 compared to the GUS control, with the strongest and most consistent increase upon expression with AtSIZ1 (Fig 8a, Fig S12).

**Fig. 8.**
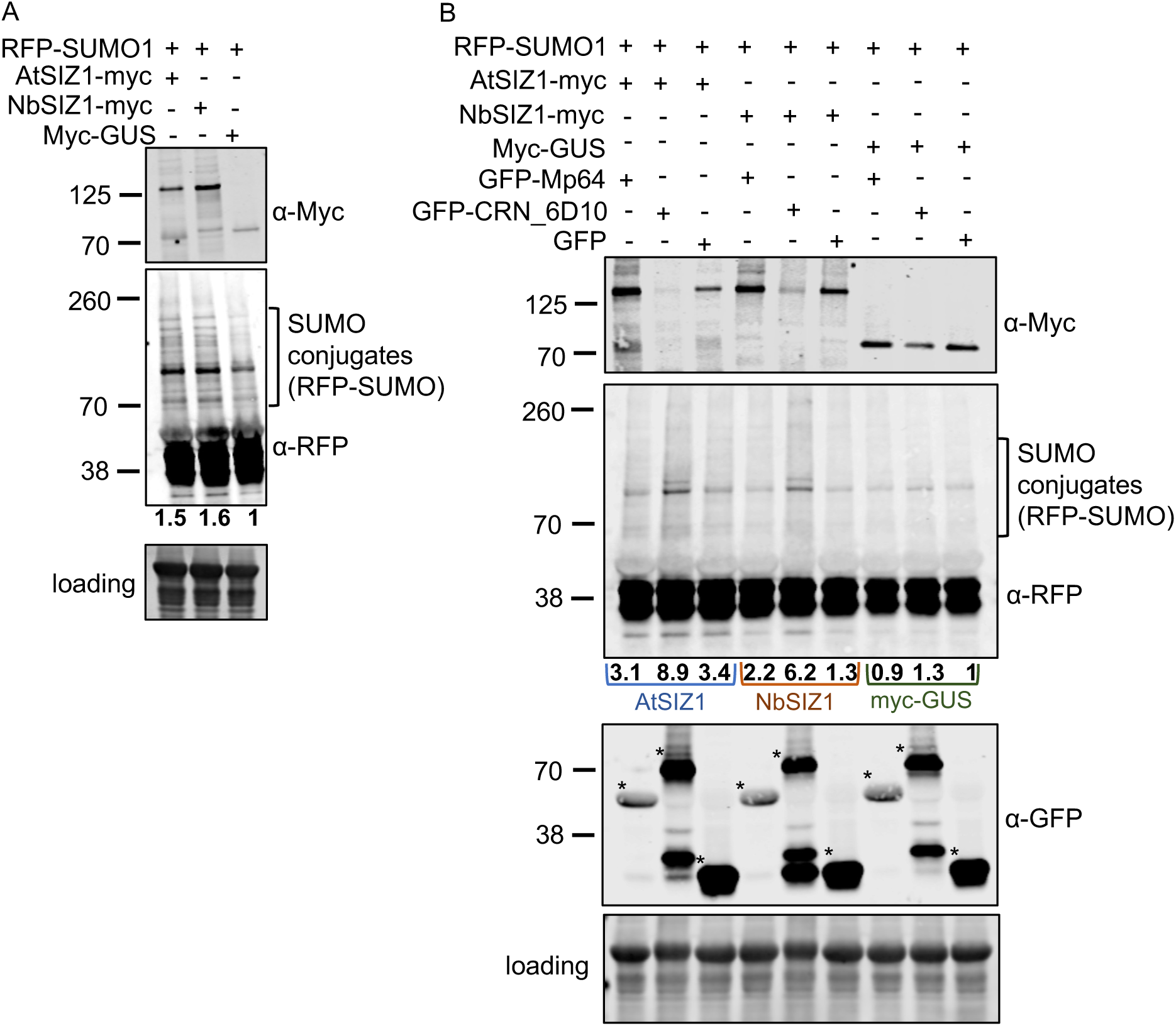
CRN83_152_6D10 enhances SIZ1-mediated SUMOylation. (A) Western blot sowing levels of SUMO-conjutages, detected using an RFP-antibody against RFP-AtSUMO1, upon ectopic/overexpression of AtSIZ1-myc, NbSIZ1-my or myc-GUS (control). Blots were prepared using total plant extracts of *N. benthamiana* infiltration sites expression different SIZ1-myc versions or myc-GUS (control). Leaf material was harvested 2pid. Total protein amount was detected using the Revert™ 700 Total Protein Stain followed by imaging in the 700 nm channel using an Odyssey^®^ CLx Imaging System. The panel indicated by “loading” shows a proportion of the membrane that includes Rubisco. Detection of SIZ1-myc fusion proteins as well as RFP-AtSUMO1 upon antibody incubation was in the 800nm channel using an Odyssey^®^ CLx Imaging System. Protein quantitation of SUMO-conjugates was done by normalizing the total band intensity of the area indicated to correspond to RFP-SUMO-conjugates to the total protein amounts using Empiria Studio 2.1 (Licor). RFP-SUMO1-conjugate levels in samples with SIZ1-myc were compared to myc-GUS (control, set at 1) to generate band intensity ratios, indicated by values below the western blot incubated with RFP-antibodies. (B) Western blot showing levels of SUMO conjugates as in (A) in the presence of GFP-Mp64, GFP-CRN83_152_6D10 or GFP (control). Western blotting and protein detection as in (A) was used to detect the presence of SIZ1-myc and GFP-effector proteins and compare levels of RFP-SUMO1-conjugates. RFP-SUMO1-conjugate levels in samples with GFP-Mp64 or GFP-CRN83_152_6D10 were compared to GFP combined with myc-GUS (control, set at 1) to generate band intensity ratios, indicated by values below the western blot incubated with RFP-antibodies.

We then performed similar co-expression assays and Western blot analyses in the presence and absence of effectors Mp64 and CRN83_152_6D10. As described above, we observed increased levels of SIZ1 in the presence of Mp64, as well as a slight decrease in the presence of CRN83_152_6D10 (Fig. 8b). Furthermore, in the presence of both SIZ1 and CRN83_152_6D10 we noted an increase in SUMO profile levels compared to the no effector (GFP) control (Fig 8b, Fig S12). We detected no increase in SUMO profile levels in the presence of CRN83_152_6D10 in combination with the GUS (control) indicating that that the observed increase in SUMOylation mediated by this effector is dependent on SIZ1 over/ectopic expression. Our data suggest that the *P. capsica* effector CRN83_152_6D10 enhances SIZ1 activity, most likely to enhance host susceptibility.

## Discussion

Pathogen infection strategies involve extensive modification of host cell biology, which rely on the modulation of hubs that control plant immunity. We show that effectors from an herbivorous insect and oomycete plant pathogen target the host E3 SUMO ligase SIZ1. Our findings suggest that the virulence strategies of two plant parasites, with distinct evolutionary histories and lifestyles, convergence on an important host immune component.

We show that SIZ1 is a key target of distinct plant parasites, which is in line with a recent study on the cyst nematode *Globodera pallida,* which shows that effector GpRbp1 associates with potato SIZ1 *in planta* (Diaz-Granados *et al*., 2019). StSIZ1 emerged as a negative regulator of immunity in plant-nematode interactions (Diaz-Granados *et al*., 2019), but the signalling requirements for this immunity have not yet been reported. We propose that SIZ1 is an important regulator of susceptibility to a broad range of plant parasites, including herbivorous insects. Indeed, Arabidopsis *siz1-2* plants show reduced susceptibility not only upon pathogen infection as reported here (Fig. 3) and previously (Lee *et al*., 2007) but also upon aphid infestation. In contrast to *siz1-2* enhanced resistance to *P. syringae pv. tomato* DC3000, which is dependent on SA, EDS1, PAD4 and SNC1, we find that resistance to the aphid *M. persicae* and the oomycete *P. capsici* is largely independent from these signalling components. These results point to 1) the involvement of a yet to be identified SIZ1-dependent signalling pathway that regulates plant immunity, and/or 2) a yet to be characterized role of SIZ1 in promoting pathogen and pest susceptibility. Although PAD4 has been reported to play an important role in plant defence against *M. persicae* (Pegadaraju *et al*., 2007), in line with Lei et al. (Lei *et al*., 2014), we did not observe an enhanced susceptibility phenotype of Arabidopsis *pad4-1* in our aphid performance assays. This may be due to differences in experimental design and conditions.

A reduction of SA levels in the *NahG* line did not enhance defence against the aphid *M. persicae* (this study and previous reports (Pegadaraju *et al*., 2007; Lei *et al*., 2014), nor did this reduce nonhost resistance to the oomycete *P. capsici,* in contrast to an earlier report by Wang et al. (Wang *et al*., 2013). However, we did observe a trend towards reduced resistance of the transgenic NahG line to *P. capsici*, but this reduction was not statistically significant and may be less pronounced due to differences in experimental set-up and infection conditions compared to Wang et al (Wang *et al*., 2013). Arabidopsis defence to insect herbivores is mediated predominantly through JA-signalling, whereas defence against (hemi-)biotrophic pathogens tend to rely on SA-signalling (Howe & Jander, 2008; Pieterse *et al*., 2012). In the Arabidopsis-*M. persicae* interaction, *siz1-2* reduced susceptibility is largely independent of SA accumulation, with the *siz1-2/NahG* line being more resistant to aphids than the *NahG* control and Col-0 (Fig. 3). Therefore, and in contrast to Lee et al.(Lee *et al*., 2007), SIZ1-regulated immunity to aphids is independent of SA-signalling. Interestingly, the Arabidopsis *siz1-2* mutant features changes in cell division, cell expansion and secondary cell wall formation, including reduced secondary cell wall thickening (Miura *et al*., 2010; Liu *et al*., 2019). Aphid feeding can trigger changes in cell wall composition that are associated with defences (Rasool *et al*., 2017), and therefore changes in cell wall formation can be responsible for altered susceptibility. However, reduced cell wall thickening most likely would lead to a reduction in defence against aphids rather than an increase as observed in the *siz1-2* mutant.

With SIZ1 comprised of several conserved domain involved in different stress responses (Cheong *et al*., 2009) it is possible that Mp64 and CRN83_152 target different protein regions and functions. Arabidopsis and *N. benthamiana* SIZ1 domains include the SAP (scaffold attachment factor A/B/acinus/PIAS) domain, PINIT (proline-isoleucine-asparagine-isoleucine-threonine) domain, an SP-RING (SIZ/PIAS-RING) domain, SXS motif (serine-X-serine), and a PHD (plant homeodomain). Functional analyses, using a set of (deletion) mutants revealed that these domains contribute differently to the wide range of SIZ1 functions in both abiotic and biotic stress (Cheong *et al*., 2009). The SP-RING domain of AtSIZ1 contributes to the nuclear localisation, SUMOylation activity, as well as the regulation of SA levels and associated plant defence responses. This domain is the suggested SIZ1 target site of the nematode effector GpRbp1 to interfere with SA-mediated defences (Diaz-Granados *et al*., 2019). Our Arabidopsis-*M. persicae* interaction assays though suggest that SIZ1 may also regulate immunity/susceptibility in a SA-independent manner where other domains may play an important role. Interestingly, SIZ1-mediated SUMOylation is involved in regulating sugar signalling independent of SA (Castro *et al*., 2015; Castro *et al*., 2018), with the *siz1-2* mutant showing reduced starch levels and increased expression of starch and sucrose catabolic genes. Aphid infestation affects sugar metabolism as reflected for example by an increase in sucrose and starch in infested Arabidopsis plants (Singh *et al*., 2011). With sugars in phloem sap also being the main aphid food source, it will be interesting to further explore a possible link between the role of SIZ1 in regulating sugar signaling and host susceptibility.

Our data support a key role for SIZ1 in host susceptibility to *P. capsici* and *M. persicae* that is targeted during infection and infestation, and point to potential different mechanisms by which effectors CRN83_152 and Mp64 modulate SIZ1 function. The presence of host SIZ1 is required for infestation/infection as knock-out of *AtSIZ1* and knock-down of *NbSIZ1* result in reduced host susceptibility phenotypes. Therefore, we propose that Mp64 and CRN83_152 redirect and perhaps enhance SIZ1 function rather than inhibit its signalling activity. Indeed, we show that CRN83_152_6D10 increased SIZ1-mediated SUMOylation *in planta*, indicating that this effector modulates E3 SUMO ligase activity (Fig. 8) . On the other hand, Mp64 but not CRN83_152_6D10 enhanced stability of SIZ1 (Fig. 6). In line with these results, we found that Arabidopsis transgenic lines expressing Mp64 do not show a reduced growth phenotype similar to the *siz1-2* mutant (Fig S4). However, expression of CRN83_152 but not Mp64 in *N. benthamiana* led to an increase in *P. capsici* infection. Based on these observations, we propose that while both virulence strategies have converged onto SIZ1, their mechanisms of action are distinct. In this context, we cannot rule out that additional candidate targets of Mp64 and CRN83_152 identified in our Y2H screens (Table S1) explain our observed differences in effector virulence activities.

As an E3 SUMO ligase, SIZ1 is required for SUMOylation of a range of substrates including chromatin modifiers, coactivators, repressors, and transcription factors that are associated with biotic and abiotic stress responses (Rytz *et al*., 2018). Similar to ubiquitination, SUMOylation involves three key steps (Verma *et al*., 2018). First, the SUMO precursor is cleaved and the SUMO moiety is linked to an SUMO-activating enzyme (E1). Activated SUMO is then transferred to the SUMO-conjugating enzyme (E2), after which it is linked to target substrates with the help of SUMO-ligases (E3). In the SUMO cycle, SUMO proteases are responsible for processing of the SUMO precursor and release of SUMO from target substrates. Given that CRN83_152 enhances SIZ1 E3 SUMO ligase activity, we hypothesize that the cell death triggered by AtSIZ1 upon transient expression in *N. benthamiana* is linked to its enzyme activity. Perhaps AtSIZ1 expression in a different plant species than Arabidopsis leads to mis-targeting of substrates, and subsequent activation of cell death. Although Mp64 did not enhance the cell death triggered by AtSIZ1, this effector did increase SIZ1 protein stability. Similarly, the effector AVR3a from *P. infestans* interacts with and stabilizes the E3 ubiquitin ligase CMPG1, likely by modifying its activity, to suppress plant immunity (Bos, JIB *et al*., 2010). The mechanism underlying the stabilization of SIZ1 by Mp64 is yet unclear. However, we hypothesize that increased stability of SIZ1, which functions as an E3 SUMO ligase, leads to increased SUMOylation activity towards its substrates and will likely affect SIZ1 complex formation with other key regulators of plant immunity.

SUMOylation of target proteins plays an important role in plant immunity and is known to be targeted as part of bacterial plant pathogen infection strategies (Verma *et al*., 2018)). For example, effector XopD from *Xanthomonas campestris pv. vesicatoria* (*Xcv*) functions as a SUMO protease inside host cells to modulate host defence signalling (Hotson *et al*., 2003; Kim *et al*., 2008). SUMOylation sites are predicted in Mp64 (Fig. S5) and CRN83_152 (1 SUMO interaction motif: SVEKGAN<u>ILSVE</u>VPGCDVD; SUMOylation site: VKMLIEV<u>K</u>REVKSAS) using prediction software GPS-SUMO (Zhao *et al*., 2014). However, using similar assays with RFP-SUMO1 described in this study, we have not detected any SUMOylation forms of Mp64 and CRN83_152, suggesting that these effectors are not SUMOylation targets themselves. Overall, our data suggest that modification of host SUMOylation is a common strategy of plant parasites to enable host colonization, and that the targeting strategies have evolved independently in distinct plant-feeding organisms including herbivorous insects. A detailed analyses of changes in the SIZ1-dependent host plant SUMOylome and their impact on susceptibility and immunity is needed to understand how distinct plant parasites promote virulence through SIZ1 targeting.

## Materials and Methods

### Plants and growth conditions

*Nicotiana benthamiana* plants were grown in a greenhouse with 16h of light, at 25° during daytime.

Transgenic Arabidopsis lines *siz1-2*, *eds1-2* (backcrossed into Col-0) (Bartsch *et al*., 2006), *pad4-1* (Glazebrook et al., 1997), *snc1-11* (Yang & Hua, 2004) and *NahG* (Delaney et al., 1994), *siz1-2*/*NahG* (Lee *et al*., 2007), *siz1-2/eds1-2* (Hammoudi *et al*., 2018), *siz1-2/pad4-1* (Lee *et al*., 2007), and *siz1-2/snc1-11* (Hammoudi *et al*., 2018) were kindly provided by Dr H.A. van den Burg, The University of Amsterdam. *Arabidopsis thaliana* plants were grown in growth chambers with an 8h light/16h dark cycle at 22°/20° (day/night), with a light intensity of 100-200 µmol/m^-2^ s^-1^ and relative humidity of 60%.

### Aphid rearing and *P. capsici* growth conditions

*M. persicae* (JHI_genotype O (Thorpe *et al*., 2018)) was maintained on oil seed rape (*Brassica napus*) plants in a Perspex growth chamber, with 12h light, at 17°C and 50% relative humidity.

*P. capsici* isolate LT1534 (obtained from Kurt Lamour, University of Tennessee) was maintained on V8 agar cubes at room temperature. For zoospore collection, P. capsici LT1534 was grown on V8 agar plates at 25°.

### Plasmid construction

The coding sequence of Mp64, lacking the region encoding the N-terminal signal peptide, was amplified from *M. persicae* (JHI_genotype O) cDNA by PCR with gene-specific primers DONR-Mp64_F and DONR-Mp64_Rev (Table S2.) The amplicon was cloned into entry vector pDONR207 (Invitrogen) using Gateway cloning technology. Cloning of the *Phythophthora capsici* effector CRN83_152 and the CRN83_152_6D10 mutant was previously described (Stam *et al*., 2013a; Amaro *et al*., 2018). For *in planta* expression, both effectors were cloned into destination vector pB7WGF2 (N-terminal GFP tag) (Karimi *et al*., 2002). For yeast-two-hybrid assays, effectors were cloned into destination vector pLexA (Dual Systems Biotech) via LR reactions using Gateway technology (Invitrogen). Vector specific primers pDONR207-F, pDONR207-R, p35s-F, GFP-Nter-F, pLexA-N-F and pLexA-N-R used in plasmid construction are listed in Table S2.

An entry clone carrying *AtSIZ1* was kindly provided by Dr H.A. van den Burg, The University of Amsterdam. *NbSIZ1* (Niben101Scf04549g09015.1, Solgenomics) was amplified from *N. benthamiana* cDNA with gene-specific primers NbSIZ1-attB1 and NbSIZ1-attB2 or NbSIZ1-attB2-nostop (Table S2). Amplicons were cloned into entry vector pDONR207 (Invitrogen) using Gateway technology. For *in planta* expression, *AtSIZ1* and *NbSIZ1* were cloned into destination vectors pB7FWG2 (C-terminal GFP tag)(Karimi *et al*., 2002), pK7RWG2 (C-terminal mRFP tag, Karimi et al., 2005), and pGWB20 (C-terminal 10xMyc tag) (Nakagawa *et al*., 2007). For yeast-two-hybrid assays, AtSIZ1 and AtSIZ1 mutants were cloned into destination vector pGAD-HA (Dual Systems Biotech) via LR reactions using Gateway technology (Invitrogen). Vector specific primers pDONR207-F, pDONR207-R, p35s-F, GFP-Cter-a-R, RFP-RevSeq, pGWB-F, pGAD-HA-F2 and pGAD-HA-R2 used in plasmid construction were listed in Table S2.

An entry clone carrying AtSUMO1 was kindly provided by Dr H.A. van den Burg, The University of Amsterdam (van den Burg *et al*., 2010). For *in planta* SUMOylation assays, AtSUMO1 was cloned into destination vector pK7WGR2 (N-terminal mRFP tag, (Karimi *et al*., 2002)) by LR reactions using Gateway technology (Invitrogen). Vector specific primer p35s-F used in plasmid construction are listed in Table S2.

### Yeast-two-hybrid assays

Yeast two hybrid screening of effectors against a *N. benthamiana* library was based on the Dualsystems Y2H system (Dual Systems Biotech) following manufacturer’s instructions. Bait vectors (pLex-N) carrying effector sequences (lacking the signal peptide encoding sequence) were transformed into yeast strain NMY51. The prey library was generated in pGAD-HA from cDNA obtained from a combination of healthy leaves, leaves infected with *P. capsici*, and leaves infested with aphids. Transformants were selected on media plates lacking leucine, tryptophan, and histidine (-LWH) with addition of 2.5mM 3-amino-1,2,4-triazole (3-AT). Yeast colonies were subjected to the β-galactosidase reporter assays according to manufacturer’s instructions. The inserts of selected yeast colonies were sequenced and analysed. The Mp64/CRN83_152-NbSIZ1 interaction was validated in yeast by independent co-transformation experiments and reporter assays.

### Generation of Arabidopsis transgenic lines by floral dipping

Arabidopsis Col-0 were grown in the greenhouse under long-day conditions (16h of light) until flowering. The flowers were dipped 3 times (one-week interval) in an *Agrobacterium* GV3101 (carrying pB7WG2-Mp64 or pB7WG2) suspension of OD_600_ =0.8-2. T1 transformants were selected using 100µg/ml BASTA (Glufosinate-ammonium) spray, and T2 seed were selected on Murashige Skoog media containing 10μg/mL BASTA. Homozygous T3 plants (predicted single insertion based on 3:1 segregation in T2) were used for aphid performance experiments. Primers Mp64-int-F/Mp64-int-Rev and Mp64-qPCR-F/Mp64-qPCR-R were used to confirm the presence of Mp64 in transgenic Arabidopsis by PCR and RT-PCR respectively (Table S2).

### SIZ1 cell death assays

*Agrobacterium* GV3101 cultures carrying C-terminal RFP-tagged AtSIZ1, NbSIZ1 or GUS were infiltrated into *N. benthamiana* leaves with an OD_600_ of 0.3, together with silencing suppressor p19 (OD_600_=0.1).

For co-expression assays, mixtures of *Agrobacterium* cultures carrying N-terminal GFP tagged Mp64, CRN83_152_6D10 or GUS with cultures carrying C-terminal RFP tagged AtSIZ1, NbSIZ1 or GUS respectively were infiltrated into *N. benthamiana* leaves (OD_600_=0.3 for each construct; for p19, OD_600_=0.1). Cell death was scored 4-7 days post inoculation using a scale of 0-3 based on the severity of the phenotype. Infiltration site were scored for no symptoms (score 0), chlorosis with localized cell death (score 1), less than 50% of the site showing visible cell death (score 2), over 50% of the infiltration site showing cell death (score 3).

Statistical analyses were conducted by using R studio Version 1.2.5001 running R-3.6.1. Differences between treatments were analysed using the Kruskal-Wallis test with post hoc Dunn’s test for multiple comparisons.

### Pathogen and pest infection/infestation assays on Arabidopsis

Two 2-day old *M. persicae* nymphs (age-synchronized) were placed on 4-6-week old Arabidopsis plants. The plants were placed in a large plastic tube sealed with a mesh lid and placed in a growth cabinet (8h of light, 22/20° for day/night, 60% humidity). Aphids were counted 10 days post infestation.

*P. capsici* isolate LT1534 was grown in V8 agar plate for 3 days in the dark at 25° and exposed to continuous light for 2 days to stimulate sporulation. Sporangia were collected in ice-cold water and incubated under light for 30-45min to promote zoospore release. For Arabidopsis infection, 4-6 weeks old plants were spray-inoculated with 100,000 spores/ml. The percentage of infected leaves was scored 8 days after inoculation.

Statistical analyses were carried out using R studio Version 1.2.5001 running R-3.6.1. A linear mixed effects model, with experimental block and biological replicate incorporated as random factors, was used for aphid fecundity assays. A linear mixed effects model, with biological replicates as a random factor, was used for *P. capsici* infection assays. ANOVA was used to analyse the final models, by using emmeans package calculating the Least Squares Means as a post hoc test.

### Infection assays on *N. benthamiana* leaves transiently expressing effectors

*Phytophthora capsici* infection assays were performed on *N. benthamiana* leaves expressing CRN83_152_6D10, Mp64 or the vector control upon agroinfiltration (OD_600_=0.3 each). Two days after infiltration, leaves were drop inoculated with 5 µL of zoospore solution (50,000 spores/mL) from strain LT1534. Lesion diameters were measured at 2 days post-inoculation.

### Virus-induced gene silencing assays

Tobacco rattle virus (TRV)-based virus-induced gene silencing (VIGS) was used to silence *NbSIZ1* in *N. benthamiana*. The VIGS construct was generated by cloning a 249-bp fragment of *NbSIZ1*, amplified with primers Sumo_Vigs_Phusion_Frag3_F and Sumo_Vigs_Phusion_Frag3_R (Table S2). To generate a TRV control, a GFP fragment was amplified using the primers eGFP_Fw and eGFP_Rv (Table S2). Amplified fragments were cloned into the TRV vector (pTRV2) (Lu *et al*., 2003) using the In-Fusion HD cloning kit (Clontech). *Agrobacterium* strains containing desired pTRV2 constructs were co-infiltrated with strains carrying pTRV1 at OD_600_=0.5 into *N. benthamiana* plants. Three weeks post infiltration, leaves at the same position of different plants were detached for quantification of *NbSIZ1* transcripts by qRT-PCR and *P.capsici* infection assays. Six independent plants were used for each VIGS construct in each replicated experiment, with a total of three replicated experiments. For infection assays, leaves were drop-inoculated with 5μL of zoospore suspension (50,000 spores per mL) of *P. capsici* strain LT1534, or for data corresponding to Fig S7 with 10ul of zoospore suspension (100,000). Lesion diameter was recorded 2-3 days post inoculation. Data analyses was carried out by using R studio Version 1.2.5001 running R-3.6.1. Group comparison was conducted by Mann-Whitney U test for non-normally distributed data.

### Confocal microscopy

*Agrobacterium* strains carrying desired constructs were infiltrated individually or in combination in *N. benthamiana* plants with an OD_600_ of 0.1. Cells were imaged at around 36 hours post infiltration using Leica TCS SP2 AOBS and Zeiss 710 confocal microscopes with HC PL FLUOTAR 63X0.9 and HCX APO L U-V 40X0.8 water-dipping lenses. GFP was excited with 488 nm from an argon laser, and emissions were detected between 500 and 530 nm. The excitation wavelength for mRFP was 561 nm and emissions were collected from 600 to 630 nm. *N. benthamiana* Histone (H2B) fused to mRFP was used as a nuclear marker (Goodin *et al*., 2007). Single optical section images or z-stacks images were collected from leaf cells those have relatively low expression level to minimize the potential artefacts. Images were projected and processed using the Image J 1.52p-Fiji (Wayne Rasband, National Institute of Health, USA).

### Detection of GFP/RFP-fusion proteins by western blotting and co-immunoprecipitation assays

*Agrobacterium* strain GV3101 expressing N-terminal GFP-tagged Mp64/CRN83_152/CRN83_152_6D10 and C-terminal 10xMyc-tagged AtSIZ1/NbSIZ1 were co-infiltrated in *N. benthamiana* leaves (OD_600_=0.3, with p19 OD_600_=0.1). Leaf samples were harvested 48 hours later. For detection of GFP and RFP fusion proteins used in localization experiments, Laemmli loading buffer (addition of 10mM DTT) was directly added to ground leaf samples followed by SDS-PAGE and western blotting. For CO-IP, equal amounts of plant material (12 leaf discs of 1.5 cm diameter) were extracted in 2ml GTEN buffer (10% glycerol, 25 mM Tris-HCl PH=7.5, 150 mM NaCl, 1mM EDTA) supplemented with protease inhibitor (S8820, Sigma-Aldrich), 2% PVPP, 0.1% NP-40 detergent and fresh 10mM DTT. Samples were incubated on ice for 10 min. The lysate was centrifuged at 14460g for 3 times, 4min/each time and supernatants were subjected to CO-IP with GFP-Trap®-M magnetic beads (Chromotek). Western blotting was performed with a monoclonal GFP antibody (Sigma-Aldrich, cat. no. G1546) and a monoclonal cMyc antibody (both at 1:3000 dilution; Santa Cruz, cat. no. SC-40) followed by anti-mouse Ig-HRP antibody (1:5000 dilution; Sigma-Aldrich, cat. no. A9044) and blots were incubated with SuperSignal Femto substrate (Thermo Scientific^TM^) and exposed to X-ray film for chemiluminescence detection.

### Detection of SUMO conjugates and altered SIZ1 stability by western blotting

To determine whether ectopic/over-expression of SIZ1 alone or in combination with effectors altered SUMOylation, we made use of a plant expression construct expression RFP-AtSUMO1 (van den Burg et al., 2010). *Agrobacterium* GV3101 strains carrying constructs to express AtSIZ1-myc/NbSIZ1-myc or myc-GUS (control) with or without GFP-Mp64/GPF-CRN83_152_6D10 or GFP (control) were combined with strains expressing RFP-AtSUMO1 for infiltration of *N*. *benthamiana* leaves. An OD_600_ of 0.3 was used for each construct, with the addition of an *Agrobacterium* strain expressing the silencing suppressor p19 (OD_600_=0.1). Forty-eight hours post infiltration, the *N. benthamiana* plants exposed to heat stress by placing them in a 37°C incubator for 1 hour.

For detection of SIZ1 protein levels in the absence/presence of effectors, side-by-side infiltrations were performed as above but without the presence of RFP-AtSUMO1.

Protein was extracted from two leaf discs (1.5 cm diameter) in 200µl GTEN buffer as described above. The protein lysate was mixed with 4x protein loading buffer (Licor, 928-40004) (with addition of 100mM DTT) by a ratio of 3:1. For each sample, 5µl was loaded into 4–20% Mini-PROTEAN^®^ TGX Gels (BioRad, 4561096), followed by SDS-PAGE (20mA/each gel, run for about 1 hour) after denaturation at 65°C for 5 minutes. The protein was subsequently transferred to PVDF membranes for 90 mins at 90V using a wet transfer system. After transfer, the membranes were stained using Revert™ 700 Total Protein Stain (Licor, 926-11021) following the manufacturer’s manual for detection of total amounts of protein. The membranes were immediately imaged in the 700 nm channel using an Odyssey^®^ CLx Imaging System.

For subsequent detection of epitope tagged proteins, membranes were blocked with Intercept® (PBS) Blocking Buffer (Licor, 927-70001) for an hour at room temperature, following an hour of primary antibody incubation in Intercept® (PBS) Blocking Buffer with 0.2% Tween® 20 (Sigma, P1379) at room temperature. After washing with PBS buffer (3 times, 5min/each), the membranes were incubated with IRDye secondary antibody in Intercept® (PBS) Blocking Buffer with addition of 0.02% SDS and 0.2% Tween® 20 (Sigma, P1379) for an hour at room temperature. After three washes with PBS buffer, the target proteins were detected in the 800 nm channel with an Odyssey^®^ CLx Imaging System.

For detection of RFP-SUMO1 and SUMO1 conjugates, a monoclonal RFP antibody raised in Rat (Chromotek, 5F8), followed by IRDye^®^ 800CW goat anti-Rat IgG secondary antibody (Licor, 926-32219) was used at a dilution of 1:3000 and 1:10000 respectively; For detection of GFP-effectors and SIZ1-myc, a monoclonal GFP antibody raised in mouse (Sigma-Aldrich, cat. no. G1546) and a monoclonal cMyc antibody raised in mouse (Santa Cruz, cat. no. SC-40) were used at a dilution of 1:3000 respectively, followed by IRDye^®^ 800CW goat anti-mouse IgG secondary antibody (Licor, 926-32210) at 1:10000 dilution.

Protein quantification was done by normalising the band intensity of SIZ1 against the total protein amounts for the SIZ1 stability assays using Empiria Studio 2.1. Quantification of SUMO conjugates was done by normalising the signal intensity of the selected MW area of the blot corresponding to SUMO conjugates against the total protein amounts. Relative ratios of signal intensity within experimental set-ups we calculated based on comparisons to relevant control samples (eg GFP, myc-GUS).

### RNA extraction and Quantitative RT-PCR

Total RNA was extracted by using RNeasy Mini Kit (Qiagen) and DNase I treated (Invitrogen™). 1 µg RNA was reverse-transcribed using SuperScript III reverse transcriptase (Sigma-Aldrich, UK) following the manufacturer’s protocol. RT-qPCR was designed following the MIQE guidelines (Bustin *et al*., 2009) with gene specific primers (Table S2). EF1α (accession number: TC19582 (At5g60390)) and PP2A (accession number: TC21939 (At1g13320)) (Liu *et al*., 2012) were used as reference genes in *N. benthamiana* and PEX4 (or UBC; accession number: AT5G25760) in *A. thaliana* (Dekkers et al. 2012).. Each 12.5 μl reaction contained 1x GoTaq® qPCR Master Mix, 1μM of each primer, 1.4 mM MgCl2, 2.4 μM CXR reference dye and a cDNA quantity of approx. 25 ng. The PCR program was set on a StepOne™ Real-Time PCR Machine (Applied Biosystems, UK) as follows: 95° for 15 mins followed by 40 cycles of 15s at 95°, 30s at 60°, and 30s at 72°. A melting curve was generated at the end of the PCR program and 2^-ΔΔCt^ value (Livak & Schmittgen, 2001) was calculated to determine the relative expression of *NbSIZ1*. Three technical replicates were performed in each run and three biological replicates were carried out.

## Acknowledgements

We thank Dr. Daniel Leybourne for advice on statistical analyses, Dr. Petra Boevink for advice on confocal microscopy, and Dr. Harrold van den Burg for providing the various Arabidopsis *siz1-2* mutant lines. We also thank Dr. Nicholas Birch (The James Hutton Institute) for allowing us to use the EPG equipment. This work was supported by the Biotechnology and Biological Sciences Research Council (grant no. BB/J005258/1 to JIBB), the European Research Council (grant no. APHIDHOST-310190 to JIBB and 310901_RETRaIN to EH), and funding from the China Scholarship Council (CSC to SL). *Phytophthora capsici* cultures were held at The James Hutton Institute under license PH\6\2015. The authors declare no conflict of interest.

## Author contributions

JB and EH conceived the study, SL, CL, TA, PR, JB and EH designed the research, SL, CL, TA, PR, and JB performed the experiments, SL, CL, TA, PR, JB and EH analyzed the data, SL, JB and EH wrote the manuscript with input from all authors. CL and TA contributed equally to this work.

## Data availability statement

Data available in article and article supplementary material.

## Supplementary Information

**Table S1.** Candidate interactors of Mp64 and CRN83_152 identified in yeast-two-hybrid screens against a *Nicotiana benthamiana* prey library.

| Effector | Protein candidate Targets | Hits | Pfam domains |
| --- | --- | --- | --- |
| Mp64 | ref\|XP_009617749.1\|PREDICTED: E3 SUMO-protein ligase SIZ1 isoform X3 [*Nicotiana tomentosiformis*] | 2 | SAP domain (IPR003034)<br>Zinc finger, RING/FYVE/PHD-type (IPR013083)<br>Zinc finger, FYVE/PHD-type (IPR011011)<br>Zinc finger, PHD-type (IPR001965)<br>Zinc finger, PHD-finger (IPR019787)<br>Zinc finger, MIZ-type (IPR004181)<br>Peptidase A2A, retrovirus, catalytic (IPR001995) |
|  | ref\|XP_009773140.1\|PREDICTED: protein LIGHT-DEPENDENT SHORT HYPOCOTYLS 10-like [*Nicotiana sylvestris*] | 2 | ALOG domain (IPR006936) |
|  | ref\|XP_009773624.1\|PREDICTED: uncharacterized protein LOC104223818 [*Nicotiana sylvestris*] | 3 | NAD(P)-binding domain (IPR016040)<br>NmrA-like domain (IPR008030)<br>Galactose-binding domain-like (IPR008979)<br>NADH:ubiquinone oxidoreductase intermediate-associated protein 30 (IPR013857) |
|  | ref\|XP_009786006.1\|PREDICTED: protein LIGHT-DEPENDENT SHORT HYPOCOTYLS 10-like [*Nicotiana sylvestris*] | 3 | ALOG domain (IPR006936) |
|  | ref\|XP_009600988.1\|PREDICTED: 60S ribosomal protein L35-like [*Nicotiana tomentosiformis*] | 2 | none |
|  | ref\|XP_009790934.1\|PREDICTED: chlorophyll a-b binding protein 40, chloroplastic-like [*Nicotiana sylvestris*] | 1 | Chlorophyll a/b binding protein domain (IPR023329) |
|  | ref\|XP_009780849.1\|chloroplast stem-loop binding protein of 41 kDa b, chloroplastic [*Nicotiana sylvestris*] | 1 | NAD(P)-binding domain (IPR016040)<br>NAD-dependent epimerase/dehydratase, N-terminal domain (IPR001509) |
|  | ref\|XP_009777441.1\|PREDICTED: magnesium-protoporphyrin IX monomethyl ester [oxidative] cyclase, chloroplastic [*Nicotiana sylvestris*] | 1 | Domain<br>Ferritin-like superfamily (IPR009078)<br>Rubrerythrin (IPR003251) |
|  | ref\|XP_009776620.1\|serine/threonine-protein phosphatase 2A 65 kDa regulatory subunit A beta isoform-like [*Nicotiana sylvestris*] | 2 | Armadillo-type fold (IPR016024)<br>Armadillo-like helical (IPR011989) |
| CRN83_152 | E3 SUMO-protein ligase SIZ1 (Niben101Scf04549g09015.1) | 3 | zf-MIZ (pfam02891); SAP (pfam02037); PHD (pfam00628) |
|  | Structure-specific endonuclease subunit SLX1<br>(Niben101Scf01771g01024.1) | 2 | GIY-YIG<br>(pfam01541) |
|  | Dynein light chain 2<br>(Niben101Scf06556g00002.1) | 2 | Dynein_light<br>(pfam01221) |
|  | Zinc finger protein<br>(Niben101Scf02509g06003.1) | 2 | zf-H2C2_2<br>(pfam13465) |
|  | 4-hydroxy-3-methylbut-2-enyl diphosphate reductase<br>(Niben101Scf33689g00006.1) | 2 | LytB<br>(pfam02401) |
|  | Inactive poly [ADP-ribose] polymerase<br>(Niben101Scf02375g00013.1) | 1 | RST (pfam12174) |
|  | Uncharacterised protein<br>(jgi\|Phyca11\|103964\|e_gw1.8.750.1) | 1 |  |
|  | 14-3-3-like protein GF14<br>(Niben101Scf00477g00017.1) | 1 | 14-3-3<br>(pfam00244) |
|  | General regulatory factor 12<br>(Niben101Scf09559g02008.1) | 1 | 14-3-3<br>(pfam00244) |

**Table S2.**
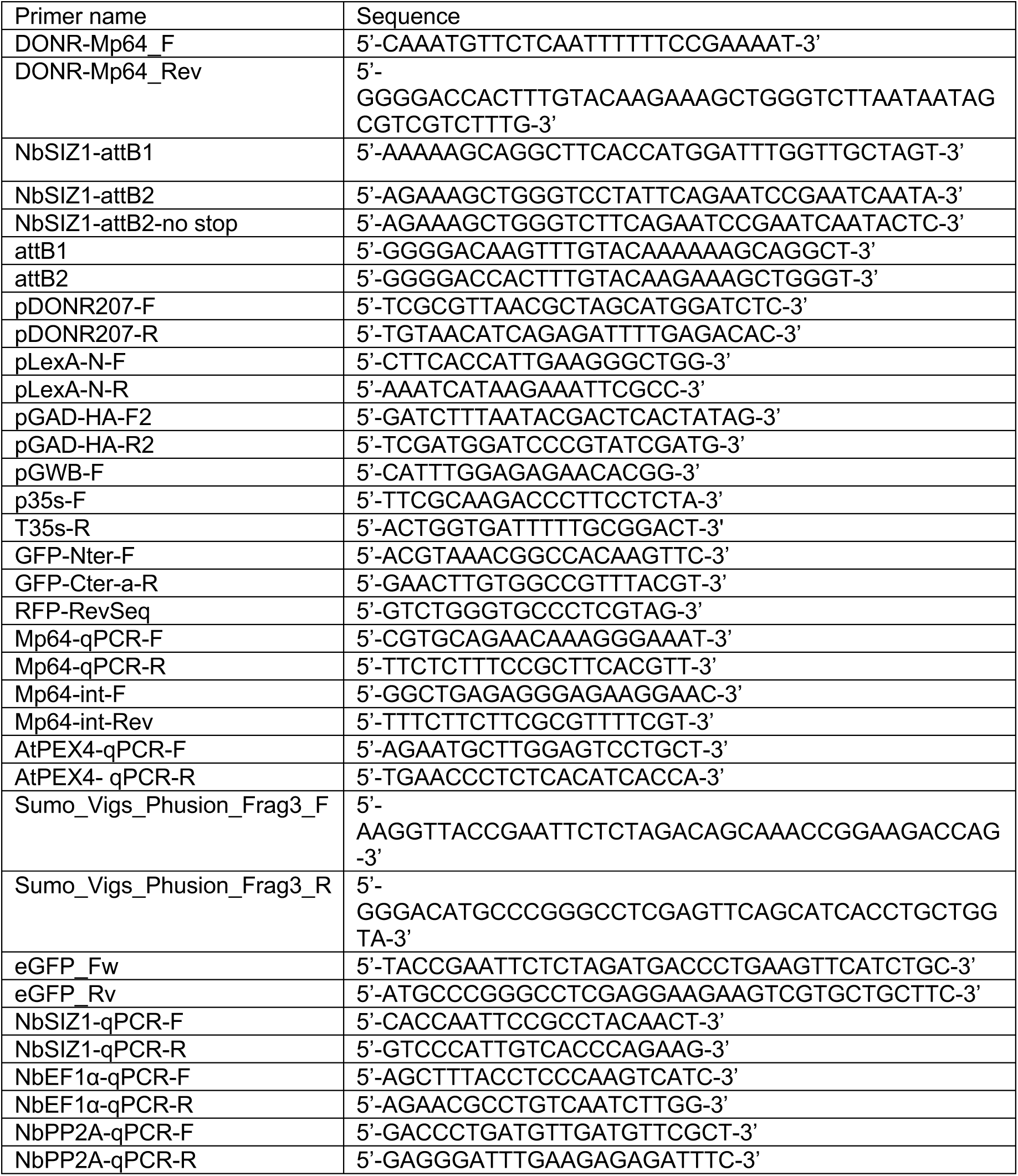
Primers used in this study.

**Table S3.** Plasmid constructs used in this study.

| Construct | Vector | Reference | Vector specific primers for cloning | Assays used |
| --- | --- | --- | --- | --- |
| LexA-Mp64 | pLexA-N | Dualsystems DUALhybrid kit | pLexA-N-F and pLexA-N-R | Yeast two hybrid |
| LexA-CRN83_152/CRN83_152_6D10 | pLexA-N | Dualsystems DUALhybrid kit | pLexA-N-F and pLexA-N-R | Yeast two hybrid |
| Untagged Mp64 | pB7WG2 | Karimi et al., 2002 | p35s-F and T35s-R | Arabidopsis transgenic line generation;<br>*Phytophthora capsici* infection assay over transient overexpression in *Nicotiana benthamiana* |
| Untagged CRN83_152 | pB7WG2 | Karimi et al., 2002 | p35s-F and T35s-R | *Phytophthora capsici* infection assay over transient overexpression in *Nicotiana benthamiana* |
| GFP-Mp64 | pB7WGF2 (N-terminal GFP tag) | Karimi et al., 2002 | p35s-F and GFP-Nter-F | CO-IP; Stability assay; SUMOylation assay; Subcellular localisation; |
| GFP-CRN83_152/CRN83_152_6D10 | pB7WGF2 (N-terminal GFP tag) | Karimi et al., 2002 | p35s-F and GFP-Nter-F | CO-IP; Stability assay; SUMOylation assay; Subcellular localisation |
| GAD-AtSIZ1 | pGAD-HA | Dualsystems DUALhybrid kit | pGAD-HA-F2 and pGAD-HA-R2 | Yeast two hybrid |
| GAD-NbSIZ1 | pGAD-HA | Dualsystems DUALhybrid kit | pGAD-HA-F2 and pGAD-HA-R2 | Yeast two hybrid |
| AtSIZ1-RFP | pK7RWG2 (C-terminal mRFP tag) | Karimi et al., 2005 | p35s-F and RFP-RevSeq | Cell death assay; Subcellular localisation |
| NbSIZ1-RFP | pK7RWG2 (C-terminal mRFP tag) | Karimi et al., 2005 | p35s-F and RFP-RevSeq | Cell death assay; Subcellular localisation |
| RFP-SUMO1 | pK7WGR2 (N-terminal mRFP tag) | Karimi et al., 2002 | p35s-F | SUMOylation assay |
| AtSIZ1-myc | pGWB20 (C-terminal 10xMyc tag) | Nakagawa et al., 2007 | pGWB-F and attB2 | CO-IP; Stability assay; SUMOylation assay |
| NbSIZ1-myc | pGWB20 (C-terminal 10xMyc tag) | Nakagawa et al., 2007 | pGWB-F and attB2 | CO-IP; Stability assay; SUMOylation assay |

**Fig. S1.**
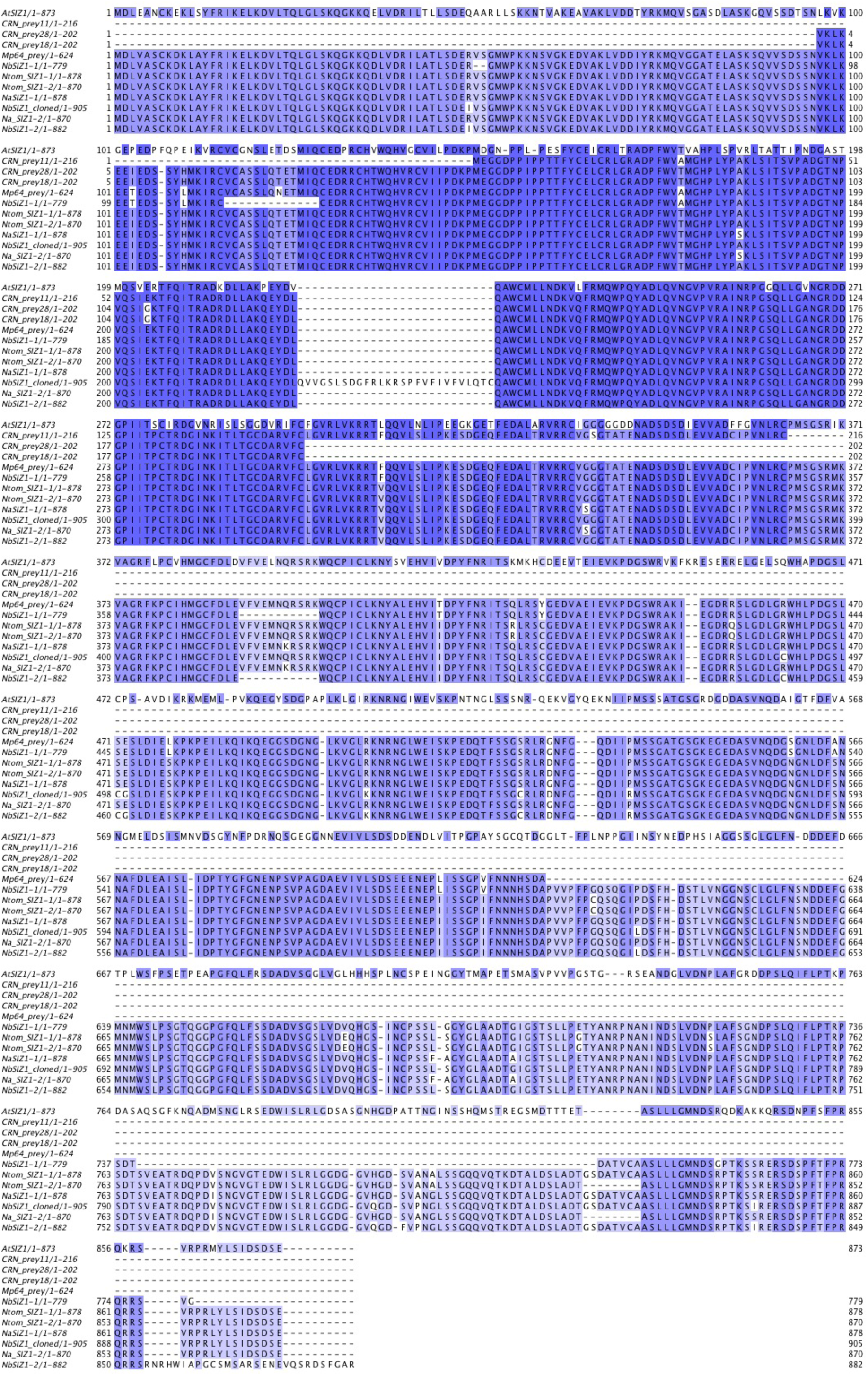
Amino acid alignment of different versions of SIZ1 and fragments recovered from yeast-two-hybrid screens. Three different prey sequences were recovered for yeast clones from the CRN83_152 screen (11, 18 and 28) and two identical prey sequences were recovered from yeast clones for the Mp64 screen. The CRN83_152 prey clones were only sequenced at the 5’-end. NbSIZ1-cloned corresponds to the NbSIZ1 sequence cloned and used in this study. AtSIZ1 corresponds to AT5G60410.1, NbSIZ1-1 corresponds to Niben101Scf15836g01010.1, NbSIZ1-2 corresponds to Niben101Scf04549g09015.1, Ntom_SIZ1-1 corresponds to XP_018631065.1, Ntom_SIZ1-2 corresponds to XP_018631066.1, NaSIZ1-1 corresponds to XP_019237907.1, NaSIZ1-2 corresponds to XP_019237903.1. Dark blue colour indicates high similarity, light blue colour low similarity.

**Fig. S2.**
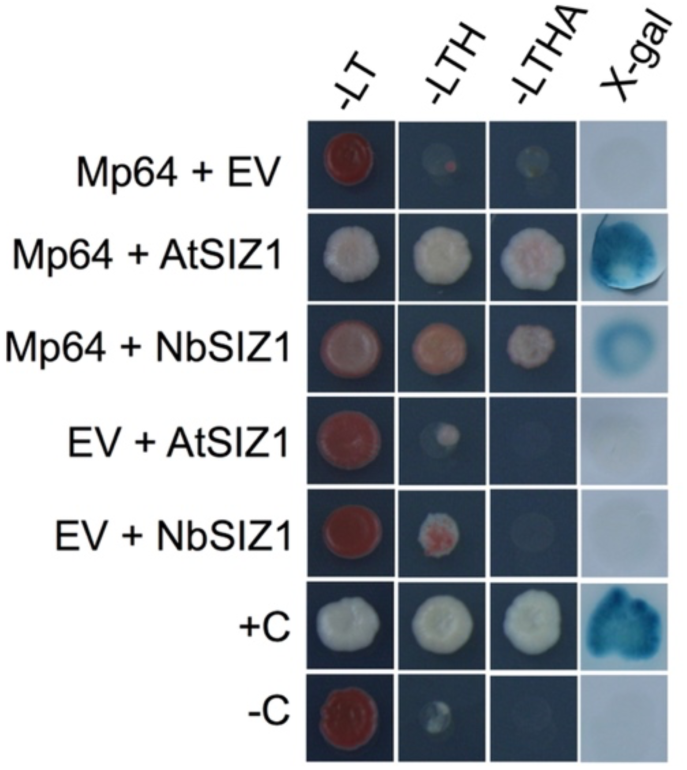
Confirmation of interactions between aphid effector Mp64 and AtSIZ1 (Arabidopsis thaliana) or NbSIZ1 (*Nicotiana benthamiana*) in yeast through activation of various reporter genes. Yeast co-transformants were selected on –LW media lacking leucine and tryptophan. -LWH represents selective medium lacking leucine, tryptophan and histidine (-LWH) and - LWHA represents selective medium lacking leucine, tryptophan, histidine and adenine. X-gal assays confirmed activation of the *lacZ* reporter gene. +C and -C indicate the positive and negative controls respectively for reporter activation, and EV indicates bait or prey vector with no insert.

**Fig. S3.**
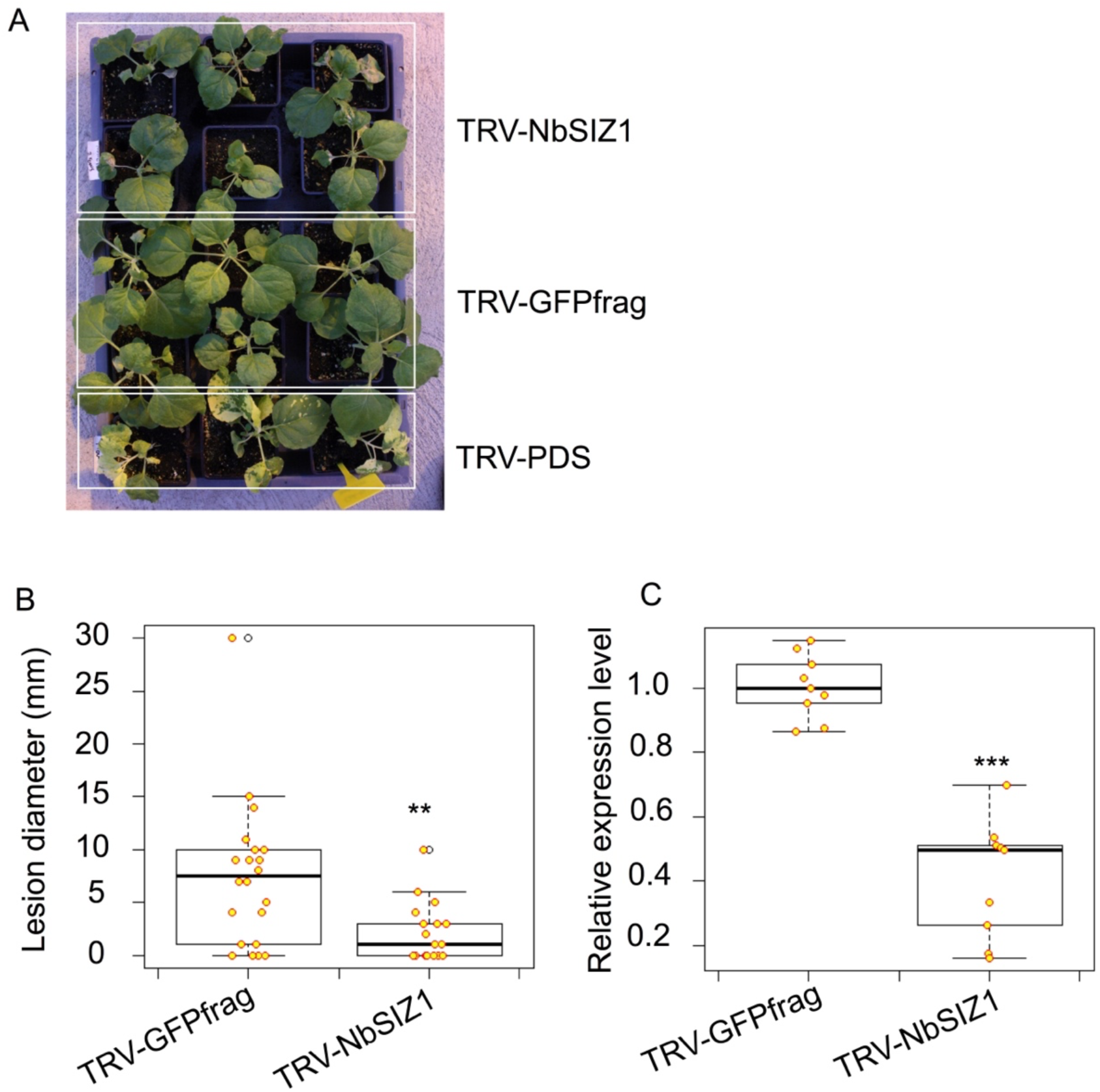
Virus-induced gene silencing of *NbSIZ1* is effective and reduces host susceptibility to *Phytophthora capsici*. (A) Representative images showing the phenotype of plants expressing TRV-*NbSIZ1* and TRV-*GFPfrag*. *NbSIZ1*-silenced plants (TRV-*NbSIZ1*) shows a slightly reduced growth and increased cell death associated virus infection compared to the control (TRV-*GFPfrag*). TRV-P*DS* was infiltrated alongside as a positive control. (B), Lesion diameter was significantly reduced in plants expressing TRV-*NbSIZ1* compared to TRV-*GFPfrag* control. Two biological replicates (n=12 per replicate) were combined in the dataset. *N. benthamiana* leaves were inoculated with 10µl of a *P. capsici* zoospores suspension (100,000 spores/ml). The lesion diameter was recorded four days post inoculation. Asterisks denote significant difference between TRV-*NbSIZ1* and TRV-*GFPfrag* (Mann-Whitney U test, p<0.001). (C) Boxplot showing that the relative expression level of *NbSIZ1* is significantly reduced in *NbSIZ1-*silenced plants (TRV-*NbSIZ1*) compared to control plants (TRV-*GFPfrag*). RT-qPCR was conducted with three biological replicates. Expression level of *NbSIZ1* was normalized against the expression of reference genes *NbPP2A* and *NbEF1α* using ΔΔCt analysis. Asterisks denote significant difference between treatments and control (Mann-Whitney U test, p<0.0001).

**Fig. S4.**
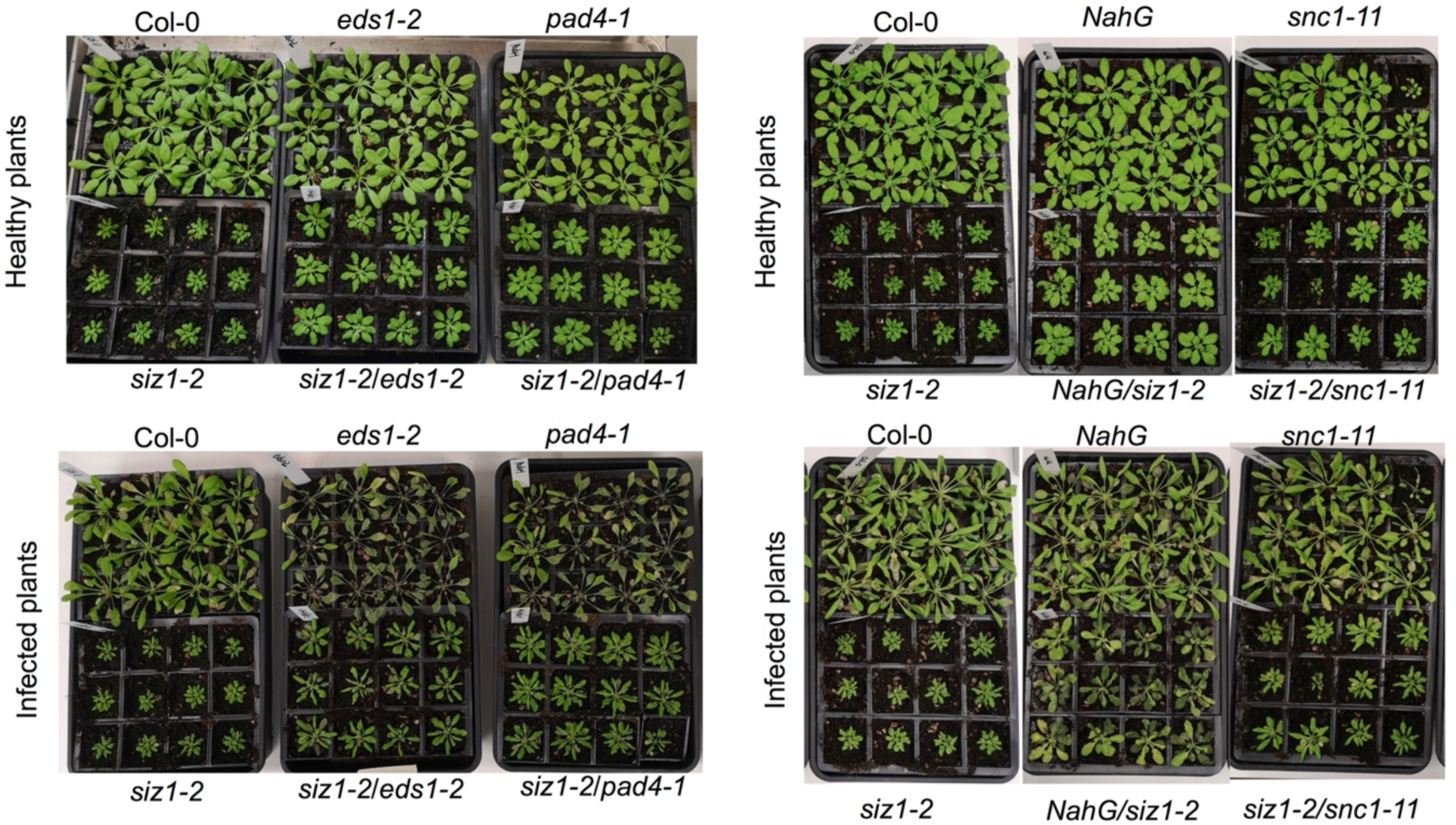
Representative images of Arabidopsis mutant lines infected with *Phytophthora capsici*. The plants were spray-inoculated with zoospore suspension of 100,000 spores/ml and the images were taken 8 days post infection.

**Fig. S5.**
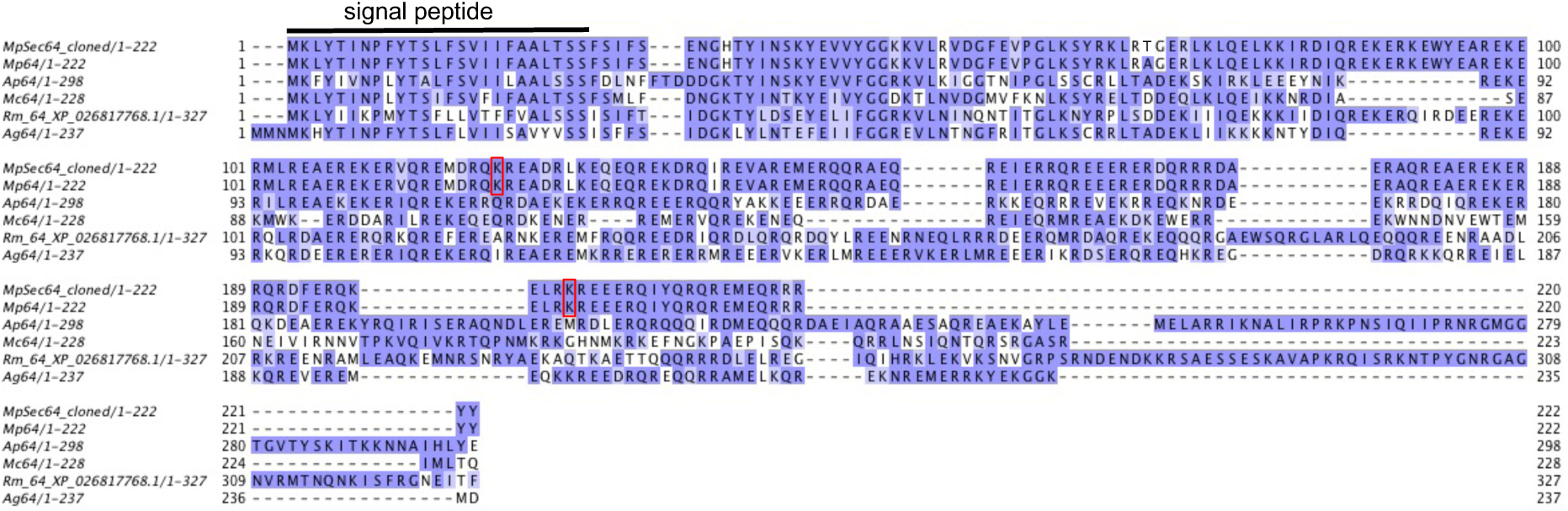
Amino acid alignment of Mp64 and predicted orthologs in aphid species. Mp64-cloned indicates the cloned effector sequence used in this study. Mp64 indicated the *Myzus persicae* database sequences XP_022180025.1. Ap64 indicates the *Acyrthosiphon pisum* sequence XP_008179242.1. Rm64 indicates the *Rhopalosiphum maidis* sequence XP_026817768.1, and Ag64 indicates the *Aphis glycines* sequence KAE9543816.1. Dark blue colour indicates high similarity, light blue colour low similarity. Signal peptide sequence of Mp64 is indicated by a black line. Red boxes indicate predicted SUMOylation sites based on GPS-SUMO 2.0 predictions with medium threshold setting.

**Fig. S6.**
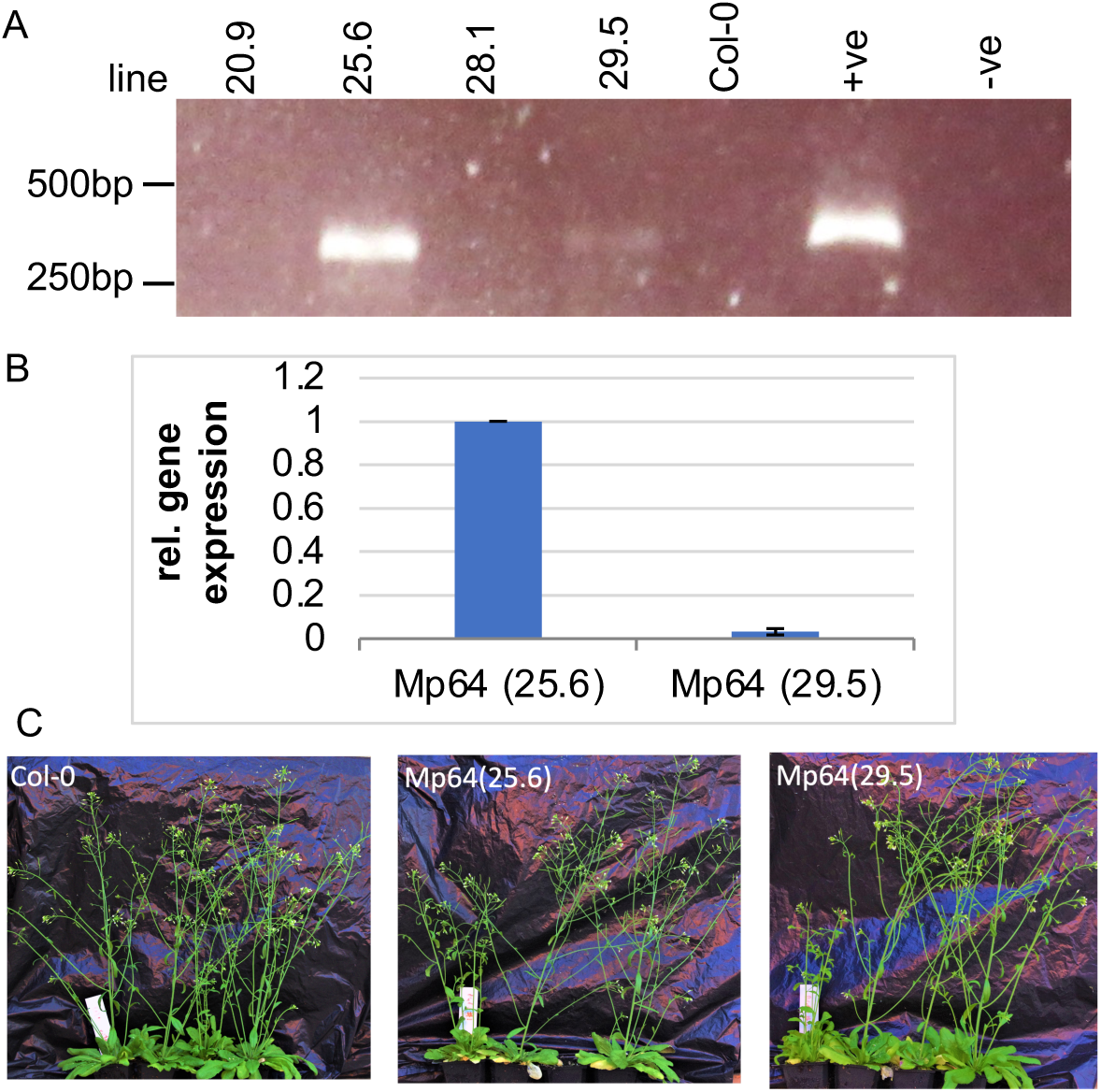
Reverse transcriptase-Polymerase Chain Reaction (RT-PCR) confirms expression of aphid effector Mp64 in transgenic Arabidopsis lines and plant phenotypes. (A) cDNA from four independent *Arabidopsis thaliana* overexpressing Mp64 lines (20.9, 25.6, 28.11, and 29.5) was used as a template for PCR with primers specific to Mp64 alongside with cDNA from Col-0, plasmid DNA with an Mp64 insert (positive control) and DNAse/RNAse treated water (negative control). The samples were analysed on a 1% agarose gel. The products from the cDNA template and plasmid were detected at the correct size of 274 bp. (B) Relative expression level of Mp64 in two independent *Arabidopsis thaliana* overexpressing line 25.6 and 29.5. RT-qPCR was conducted with three technical replicates. Expression level of Mp64 was normalised against the expression of a reference gene *AtPEX4* using ΔΔCt analysis. (C) Representative images showing the phenotype of two selected Arabidopsis Mp64 transgenic lines (line 25.6 and line 29.5).

**Fig. S7.**
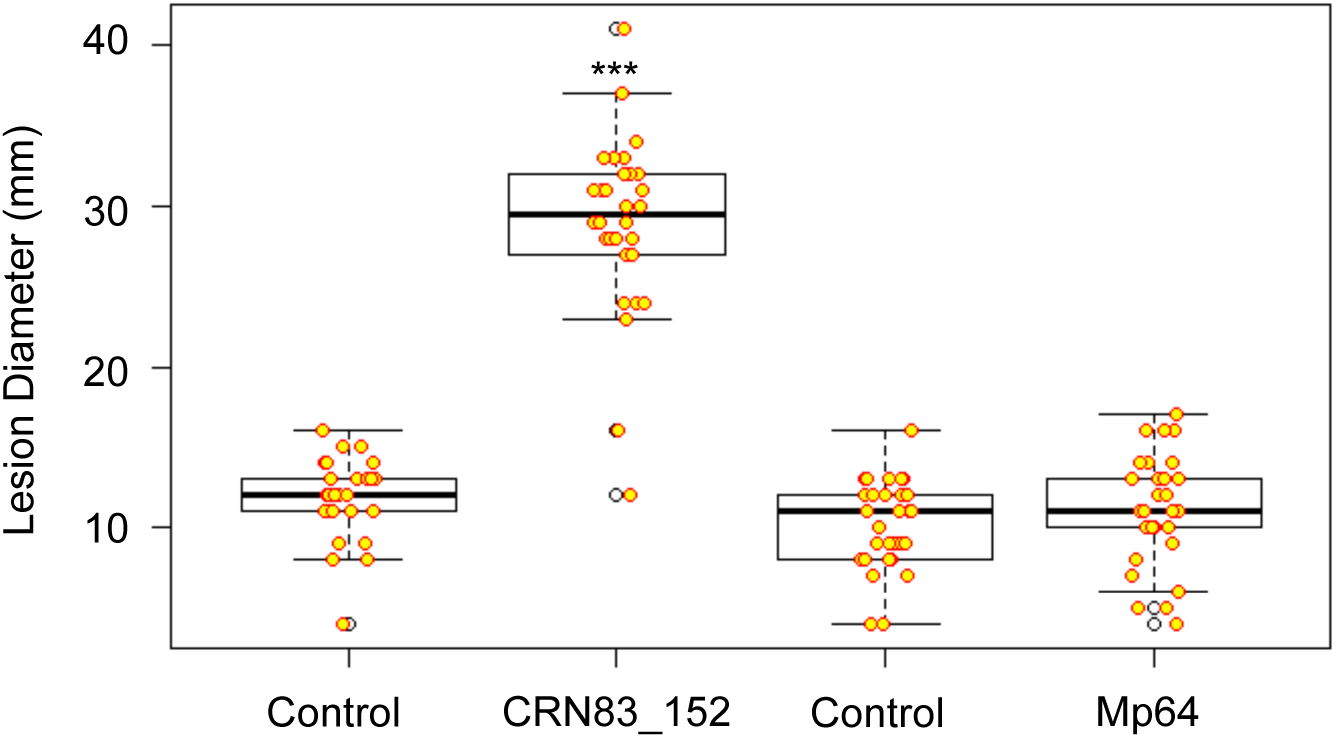
Expression of CRN83_152 but not Mp64 in *Nicotiana benthamiana* enhances host susceptibility to *Phytophthora capsici*. Leaves expressing CRN83_152, Mp64 and vector controls upon agroinfiltration were challenged with 5µl *P. capsici* zoospore suspension (50,000 spores/mL) and the lesion diameter was measured two days post inoculation. A total of 30 infiltration sites per treatment were analyzed (n=30). Asterisks denote significant difference of lesion diameter between the CRN83_152 overexpressing leaves and vector control (Mann-Whitney U test, p<0.05).

**Fig. S8.**
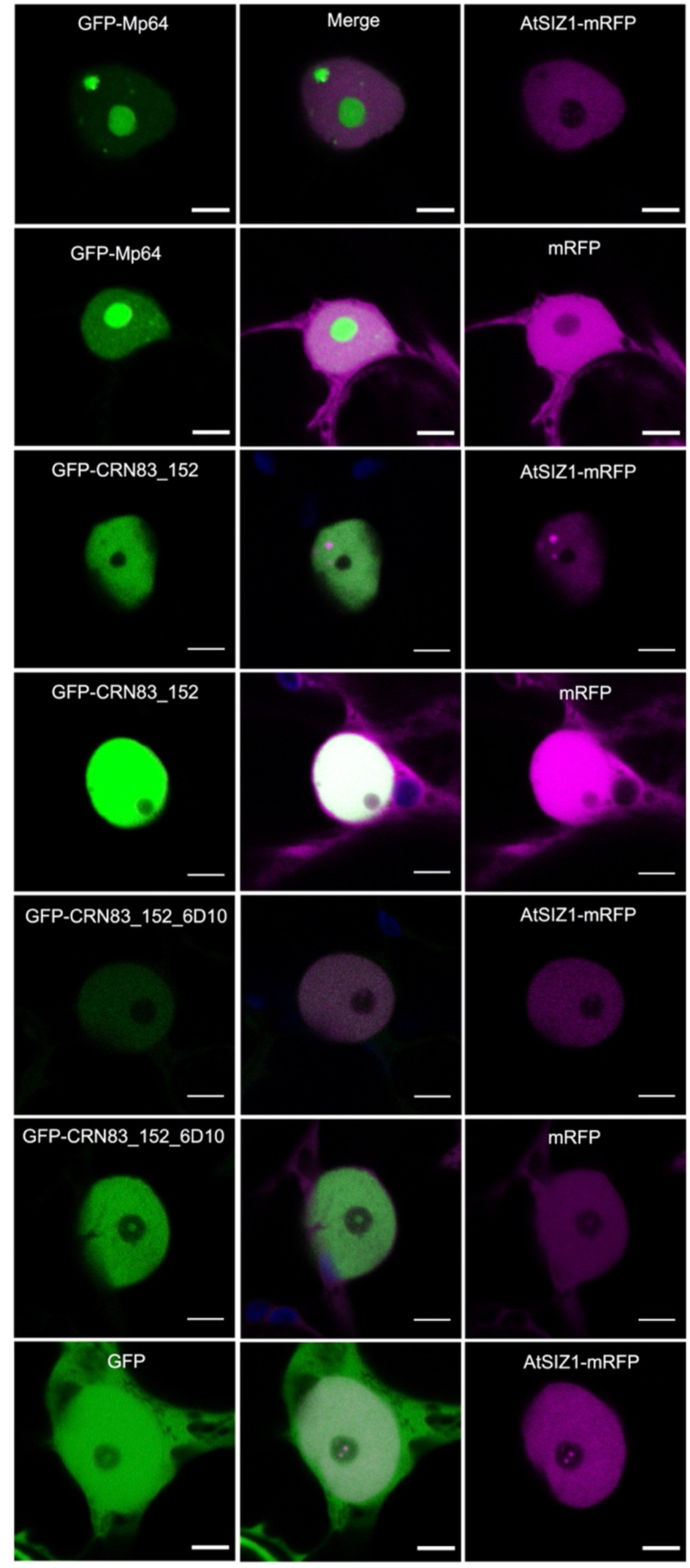
Localisation of effectors Mp64 and CRN83_152(6D1) and AtSIZ1 in the host nucleus. Leaves transiently expressing GFP-Mp64, GFP-CRN83_152 or GFP-CRN83_152_6D10 in combination with RFP or AtSIZ1 –RFP were used for confocal imaging around 36 hours after agroinfiltration. Images show single optical sections through nuclei co-expressing the GFP-effector with AtSIZ1-RFP or RFP control. Scale bars represent 5 µm.

**Fig. S9.**
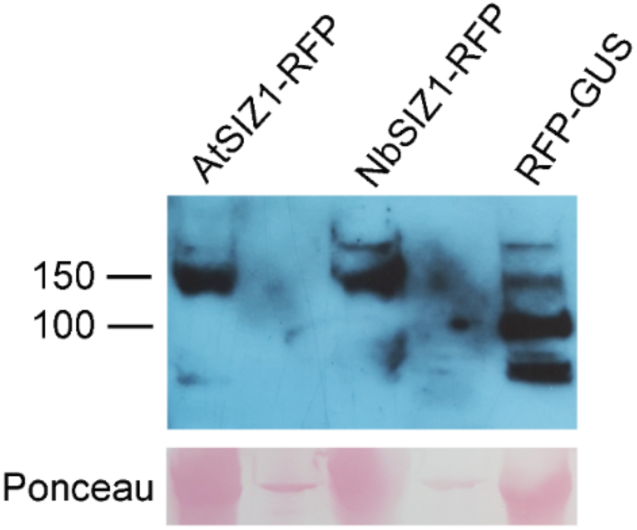
Expression of full length SIZ1-RFP fusion proteins *in planta*. Western blots showing SIZ1-RFP proteins are expressed as full-length fusion proteins. Leaf samples from infiltration sites in N. benthamiana were collected 3 days after infiltration. Leaf samples were ground in sample buffer and equal amounts were loaded on an SDS-PAGE gel for western blotting with an RFP-antibody. Marker indicated molecular weight in kD. Ponceau staining shows equal loading and transfer.

**Fig. S10.**
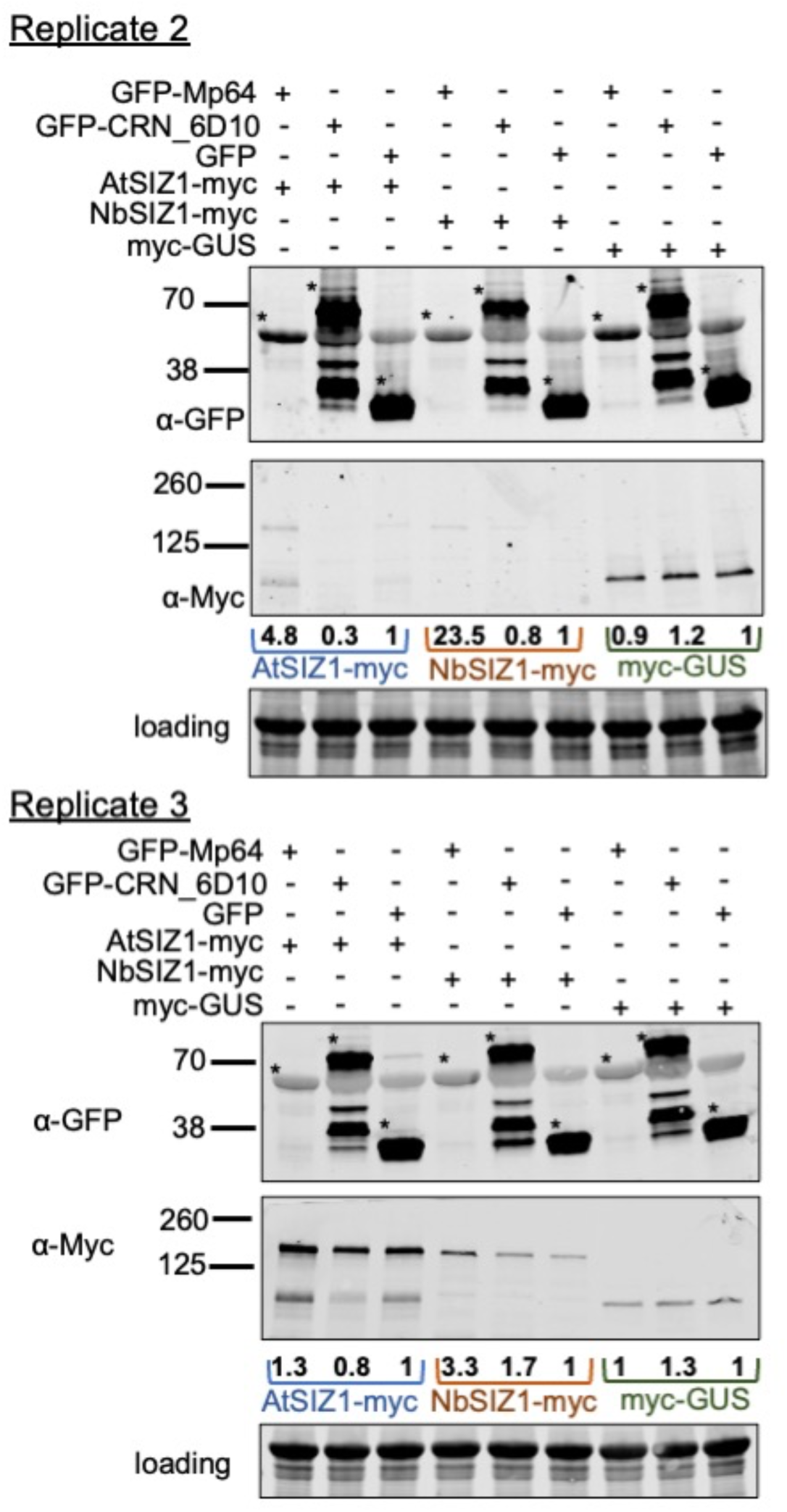
Mp64 but not CRN83-152_6D10 enhanced SIZ1 stability *in planta.* Additional replicates of western blots shown in Fig. 6. Western blots showing Mp64 increases SIZ1 protein levels. Blots were prepared using total plant extracts of *N. benthamiana* infiltration sites expression GFP-Mp64/CRN83-152_D610 or GFP (control) with SIZ1-myc. Leaf material was harvested 2 dpi. Total protein amount was detected using the Revert™ 700 Total Protein Stain followed by imaging in the 700 nm channel using an Odyssey^®^ CLx Imaging System. The panel indicated by “loading” shows a proportion of the membrane that includes Rubisco. Detection of GFP-fusion and SIZ1-myc fusion proteins upon antibody incubation was in the 800nm channel using an Odyssey^®^ CLx Imaging System. Asterisk indicated bands corresponding to GFP-effectors/GFP. SIZ1 protein quantitation was done by normalizing the band intensity of SIZ1-myc to the total protein amounts using Empiria Studio 2.1 (Licor). SIZ1-myc levels in samples with GFP-Mp64 or GFP-CRN83_152_6D10 were compared to GFP (control, set at 1) to generate band intensity ratios, indicated by values below the western blot incubated with Myc-antibodies.

**Fig. S11.**
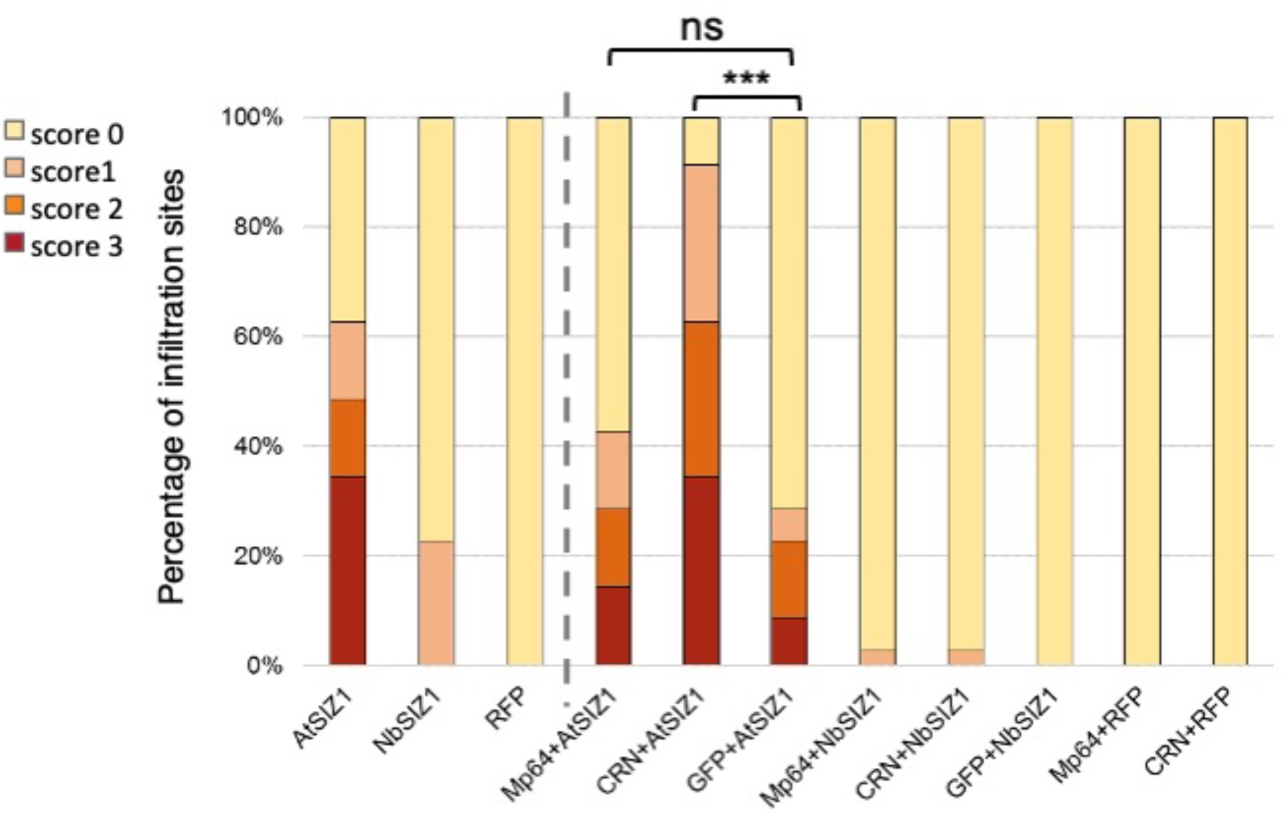
CRN83_152_6D10 enhances AtSIZ1 triggered cell death in *Nicotiana benthamiana*. Bar graph showing the proportions of infiltration sites with different levels of cell death upon expression of SIZ1 (with C-terminal RFP tag) either alone or in combination with aphid effector Mp64 or *Phytophthora capsica* effector CRN83_152_6D10 (both effectors with GFP tag), or a GFP control. Data was collected 4 days after infiltration. Graph represent data from a combination of 2 biological replicates of 11-12 infiltration sites per experiment (n=23). Note that the third biological replicate, not included here, did not yet show cell death symptoms at 4pdi. *** indicates p < 0.001 (Kruskal-Wallis test with post-hoc Dunn’s test for multiple comparison,) and ns indicates no significant difference.

**Fig. S12.**
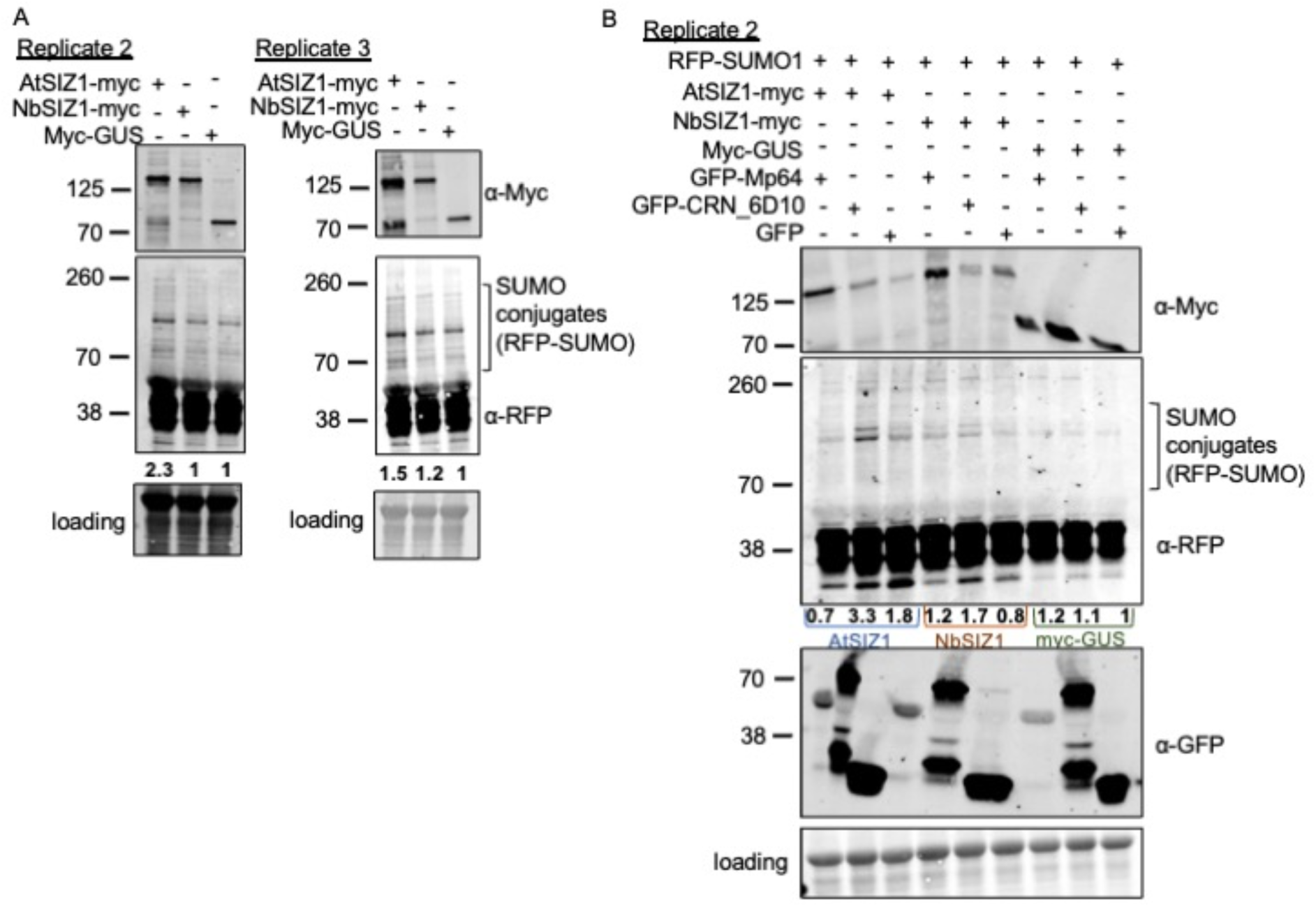
CRN83_152_6D10 enhances SIZ1-mediated SUMOylation. (A) Western blot sowing levels of SUMO-conjugates, detected using an RFP-antibody against RFP-AtSUMO1, upon ectopic/overexpression of AtSIZ1-myc, NbSIZ1-myc or myc-GUS (control). Blots were prepared using total plant extracts of *N. benthamiana* infiltration sites expression different SIZ1-myc versions or GUS-myc (control). Leaf material was harvested 2 dpi. Total protein amount was detected using the Revert™ 700 Total Protein Stain followed by imaging in the 700 nm channel using an Odyssey^®^ CLx Imaging System. The panel indicated by “loading” shows a proportion of the membrane that includes Rubisco. Detection of SIZ1-myc fusion proteins as well as RFP-AtSUMO1 upon antibody incubation was in the 800nm channel using an Odyssey^®^ CLx Imaging System. Quantification of SUMO-conjugates was done by normalizing the total band intensity of the area indicated to correspond to RFP-SUMO-conjugates to the total protein amounts using Empiria Studio 2.1 (Licor). RFP-SUMO1-conjugate levels in samples with SIZ1-myc were compared to myc-GUS(control, set at 1) to generate band intensity ratios, indicated by values below the western blot incubated with RFP-antibodies. (B) Western blot showing levels of SUMO conjugates as in (A) in the presence of GFP-Mp64, GFP-CRN83_152_6D10 or GFP (control). Western blotting and protein detection as in (A) was used to detect the presence of SIZ1-myc and GFP-effector proteins and compare levels of RFP-SUMO1-conjugates. RFP-SUMO1-conjugate levels in samples with GFP-Mp64 or GFP-CRN83_152_6D10 were compared to GFP combined with myc-GUS (control, set at 1) to generate band intensity ratios, indicated by values below the western blot incubated with RFP-antibodies.

## References

Amaro TMMM, Thilliez GJA, Mcleod RA, Huitema E. 2018. Random mutagenesis screen shows that Phytophthora capsici CRN83_152-mediated cell death is not required for its virulence function(s). Mol Plant Pathol 19(5): 1114–1126.

Bartsch M, Gobbato E, Bednarek P, Debey S, Schultze JL, Bautor J, Parker JE. 2006. Salicylic acid-independent ENHANCED DISEASE SUSCEPTIBILITY1 signaling in Arabidopsis immunity and cell death is regulated by the monooxygenase FMO1 and the Nudix hydrolase NUDT7. Plant Cell 18(4): 1038–1051.

Bos JI, Prince D, Pitino M, Maffei ME, Win J, Hogenhout SA. 2010. A functional genomics approach identifies candidate effectors from the aphid species Myzus persicae (green peach aphid). PLoS Genet 6(11): e1001216.

Bos JIB, Armstrong MR, Gilroy EM, Boevink PC, Hein I, Taylor RM, Tian ZD, Engelhardt S, Vetukuri RR, Harrower B, et al. 2010. Phytophthora infestans effector AVR3a is essential for virulence and manipulates plant immunity by stabilizing host E3 ligase CMPG1. Proceedings of the National Academy of Sciences of the United States of America 107(21): 9909–9914.

Boulain H, Legeai F, Guy E, Morlière S, Douglas NE, Oh J, Murugan M, Smith M, Jaquiéry J, Peccoud J, et al. 2018. Fast Evolution and Lineage-Specific Gene Family Expansions of Aphid Salivary Effectors Driven by Interactions with Host-Plants. Genome Biology and Evolution 10(6): 1554–1572.

Bustin SA, Benes V, Garson JA, Hellemans J, Huggett J, Kubista M, Mueller R, Nolan T, Pfaffl MW, Shipley GL, et al. 2009. The MIQE guidelines: minimum information for publication of quantitative real-time PCR experiments. Clin Chem 55(4): 611–622.

Carolan JC, Caragea D, Reardon KT, Mutti NS, Dittmer N, Pappan K, Cui F, Castaneto M, Poulain J, Dossat C, et al. 2011. Predicted effector molecules in the salivary secretome of the pea aphid (Acyrthosiphon pisum): a dual transcriptomic/proteomic approach. J Proteome Res 10(4): 1505–1518.

Carolan JC, Fitzroy CI, Ashton PD, Douglas AE, Wilkinson TL. 2009. The secreted salivary proteome of the pea aphid Acyrthosiphon pisum characterised by mass spectrometry. Proteomics 9(9): 2457–2467.

Castro PH, Verde N, Lourenço T, Magalhães AP, Tavares RM, Bejarano ER, Azevedo H. 2015. SIZ1-Dependent Post-Translational Modification by SUMO Modulates Sugar Signaling and Metabolism in Arabidopsis thaliana. Plant Cell Physiol 56(12): 2297–2311.

Castro PH, Verde N, Tavares RM, Bejarano ER, Azevedo H. 2018. Sugar signaling regulation by arabidopsis SIZ1-driven sumoylation is independent of salicylic acid. Plant Signal Behav 13(4): e1179417.

Catala R, Ouyang J, Abreu IA, Hu Y, Seo H, Zhang X, Chua NH. 2007. The Arabidopsis E3 SUMO ligase SIZ1 regulates plant growth and drought responses. Plant Cell 19(9): 2952–2966.

Chaudhary R, Peng HC, He J, MacWilliams J, Teixeira M, Tsuchiya T, Chesnais Q, Mudgett MB, Kaloshian I. 2019. Aphid effector Me10 interacts with tomato TFT7, a 14-3-3 isoform involved in aphid resistance. New Phytologist 221(3): 1518–1528.

Cheong MS, Park HC, Hong MJ, Lee J, Choi W, Jin JB, Bohnert HJ, Lee SY, Bressan RA, Yun DJ. 2009. Specific domain structures control abscisic acid-, salicylic acid-, and stress-mediated SIZ1 phenotypes. Plant Physiol 151(4): 1930–1942.

Delaney TP, Uknes S, Vernooij B, Friedrich L, Weymann K, Negrotto D, Gaffney T, Gut-Rella M, Kessmann H, Ward E, et al. 1994. A central role of salicylic Acid in plant disease resistance. Science 266(5188): 1247–1250.

Diaz-Granados A, Sterken MG, Persoon J, Overmars H, Pokhare SS, Mazur MJ, Martin-Ramirez S, Holterman M, Martin EC, Pomp R, et al. 2019. SIZ1 is a nuclear target of the nematode effector GpRbp1 from Globodera pallida that acts as a negative regulator of basal plant defense to cyst nematodes. bioRxiv.

Glazebrook J, Zook M, Mert F, Kagan I, Rogers EE, Crute IR, Holub EB, Hammerschmidt R, Ausubel FM. 1997. Phytoalexin-deficient mutants of Arabidopsis reveal that PAD4 encodes a regulatory factor and that four PAD genes contribute to downy mildew resistance. Genetics 146(1): 381–392.

Goodin MM, Chakrabarty R, Banerjee R, Yelton S, Debolt S. 2007. New gateways to discovery. Plant Physiol 145(4): 1100–1109.

Gou M, Huang Q, Qian W, Zhang Z, Jia Z, Hua J. 2017. Sumoylation E3 Ligase SIZ1 Modulates Plant Immunity Partly through the Immune Receptor Gene SNC1 in Arabidopsis. Mol Plant Microbe Interact 30(4): 334–342.

Hammoudi V, Fokkens L, Beerens B, Vlachakis G, Chatterjee S, Arroyo-Mateos M, Wackers PFK, Jonker MJ, van den Burg HA. 2018. The Arabidopsis SUMO E3 ligase SIZ1 mediates the temperature dependent trade-off between plant immunity and growth. PLoS Genet 14(1): e1007157.

Hotson A, Chosed R, Shu H, Orth K, Mudgett MB. 2003. Xanthomonas type III effector XopD targets SUMO-conjugated proteins in planta. Mol Microbiol 50(2): 377–389.

Howe GA, Jander G. 2008. Plant immunity to insect herbivores. Annu Rev Plant Biol 59: 41–66.

Huot B, Yao J, Montgomery BL, He SY. 2014. Growth-defense tradeoffs in plants: a balancing act to optimize fitness. Mol Plant 7(8): 1267–1287.

Ishida T, Yoshimura M, Miura K, Sugimoto K. 2012. MMS21/HPY2 and SIZ1, two Arabidopsis SUMO E3 ligases, have distinct functions in development. PLoS One 7(10): e46897.

Jin JB, Jin YH, Lee J, Miura K, Yoo CY, Kim WY, Van Oosten M, Hyun Y, Somers DE, Lee I, et al. 2008. The SUMO E3 ligase, AtSIZ1, regulates flowering by controlling a salicylic acid-mediated floral promotion pathway and through affects on FLC chromatin structure. Plant J 53(3): 530–540.

Jones JD, Dangl JL. 2006. The plant immune system. Nature 444(7117): 323–329.

Kaloshian I, Walling LL. 2016. Hemipteran and dipteran pests: Effectors and plant host immune regulators. J Integr Plant Biol 58(4): 350–361.

Karimi M, Inzé D, Depicker A. 2002. GATEWAY vectors for Agrobacterium-mediated plant transformation. Trends in Plant Science 7(5): 193–195.

Kim JG, Taylor KW, Hotson A, Keegan M, Schmelz EA, Mudgett MB. 2008. XopD SUMO protease affects host transcription, promotes pathogen growth, and delays symptom development in xanthomonas-infected tomato leaves. Plant Cell 20(7): 1915–1929.

Lee J, Nam J, Park HC, Na G, Miura K, Jin JB, Yoo CY, Baek D, Kim DH, Jeong JC, et al. 2007. Salicylic acid-mediated innate immunity in Arabidopsis is regulated by SIZ1 SUMO E3 ligase. Plant J 49(1): 79–90.

Lei J, A Finlayson S, Salzman RA, Shan L, Zhu-Salzman K. 2014. BOTRYTIS-INDUCED KINASE1 Modulates Arabidopsis Resistance to Green Peach Aphids via PHYTOALEXIN DEFICIENT4. Plant Physiol 165(4): 1657–1670.

Lin XL, Niu D, Hu ZL, Kim DH, Jin YH, Cai B, Liu P, Miura K, Yun DJ, Kim WY, et al. 2016. An Arabidopsis SUMO E3 Ligase, SIZ1, Negatively Regulates Photomorphogenesis by Promoting COP1 Activity. PLoS Genet 12(4): e1006016.

Liu C, Yu H, Li L. 2019. SUMO modification of LBD30 by SIZ1 regulates secondary cell wall formation in Arabidopsis thaliana. PLoS Genet 15(1): e1007928.

Liu D, Shi L, Han C, Yu J, Li D, Zhang Y. 2012. Validation of reference genes for gene expression studies in virus-infected Nicotiana benthamiana using quantitative real-time PCR. PLoS One 7(9): e46451.

Livak KJ, Schmittgen TD. 2001. Analysis of relative gene expression data using real-time quantitative PCR and the 2(-Delta Delta C(T)) Method. Methods 25(4): 402–408.

Lozano-Torres JL, Wilbers RH, Gawronski P, Boshoven JC, Finkers-Tomczak A, Cordewener JH, America AH, Overmars HA, Van ’t Klooster JW, Baranowski L, et al. 2012. Dual disease resistance mediated by the immune receptor Cf-2 in tomato requires a common virulence target of a fungus and a nematode. Proc Natl Acad Sci U S A 109(25): 10119–10124.

Lu R, Martin-Hernandez AM, Peart JR, Malcuit I, Baulcombe DC. 2003. Virus-induced gene silencing in plants. Methods 30(4): 296–303.

Mafurah JJ, Ma H, Zhang M, Xu J, He F, Ye T, Shen D, Chen Y, Rajput NA, Dou D. 2015. A Virulence Essential CRN Effector of Phytophthora capsici Suppresses Host Defense and Induces Cell Death in Plant Nucleus. PLoS One 10(5): e0127965.

Miura K, Jin JB, Lee J, Yoo CY, Stirm V, Miura T, Ashworth EN, Bressan RA, Yun DJ, Hasegawa PM. 2007. SIZ1-mediated sumoylation of ICE1 controls CBF3/DREB1A expression and freezing tolerance in Arabidopsis. Plant Cell 19(4): 1403–1414.

Miura K, Lee J, Jin JB, Yoo CY, Miura T, Hasegawa PM. 2009. Sumoylation of ABI5 by the Arabidopsis SUMO E3 ligase SIZ1 negatively regulates abscisic acid signaling. Proc Natl Acad Sci U S A 106(13): 5418–5423.

Miura K, Lee J, Miura T, Hasegawa PM. 2010. SIZ1 controls cell growth and plant development in Arabidopsis through salicylic acid. Plant Cell Physiol 51(1): 103–113.

Miura K, Rus A, Sharkhuu A, Yokoi S, Karthikeyan AS, Raghothama KG, Baek D, Koo YD, Jin JB, Bressan RA, et al. 2005. The Arabidopsis SUMO E3 ligase SIZ1 controls phosphate deficiency responses. Proc Natl Acad Sci U S A 102(21): 7760–7765.

Monaghan J, Zipfel C. 2012. Plant pattern recognition receptor complexes at the plasma membrane. Curr Opin Plant Biol 15(4): 349–357.

Mukhtar MS, Carvunis AR, Dreze M, Epple P, Steinbrenner J, Moore J, Tasan M, Galli M, Hao T, Nishimura MT, et al. 2011. Independently evolved virulence effectors converge onto hubs in a plant immune system network. Science 333(6042): 596–601.

Nakagawa T, Suzuki T, Murata S, Nakamura S, Hino T, Maeo K, Tabata R, Kawai T, Tanaka K, Niwa Y, et al. 2007. Improved Gateway binary vectors: high-performance vectors for creation of fusion constructs in transgenic analysis of plants. Biosci Biotechnol Biochem 71(8): 2095–2100.

Nguyen Ba AN, Pogoutse A, Provart N, Moses AM. 2009. NLStradamus: a simple Hidden Markov Model for nuclear localization signal prediction. BMC Bioinformatics 10: 202.

Pegadaraju V, Louis J, Singh V, Reese JC, Bautor J, Feys BJ, Cook G, Parker JE, Shah J. 2007. Phloem-based resistance to green peach aphid is controlled by Arabidopsis PHYTOALEXIN DEFICIENT4 without its signaling partner ENHANCED DISEASE SUSCEPTIBILITY1. Plant Journal 52(2): 332–341.

Pieterse CM, Van der Does D, Zamioudis C, Leon-Reyes A, Van Wees SC. 2012. Hormonal modulation of plant immunity. Annu Rev Cell Dev Biol 28: 489–521.

Rao W, Zheng X, Liu B, Guo Q, Guo J, Wu Y, Shangguan X, Wang H, Wu D, Wang Z, et al. 2019. Secretome Analysis and In Planta Expression of Salivary Proteins Identify Candidate Effectors from the Brown Planthopper Nilaparvata lugens. Mol Plant Microbe Interact 32(2): 227–239.

Rasool B, McGowan J, Pastok D, Marcus SE, Morris JA, Verrall SR, Hedley PE, Hancock RD, Foyer CH. 2017. Redox Control of Aphid Resistance through Altered Cell Wall Composition and Nutritional Quality. Plant Physiol 175(1): 259–271.

Ratcliff F, Martin-Hernandez AM, Baulcombe DC. 2001. Technical Advance. Tobacco rattle virus as a vector for analysis of gene function by silencing. Plant J 25(2): 237–245.

Rodriguez PA, Escudero-Martinez C, Bos JI. 2017. An Aphid Effector Targets Trafficking Protein VPS52 in a Host-Specific Manner to Promote Virulence. Plant Physiology 173(3): 1892–1903.

Rytz TC, Miller MJ, McLoughlin F, Augustine RC, Marshall RS, Juan YT, Charng YY, Scalf M, Smith LM, Vierstra RD. 2018. SUMOylome Profiling Reveals a Diverse Array of Nuclear Targets Modified by the SUMO Ligase SIZ1 during Heat Stress. Plant Cell 30(5): 1077–1099.

Singh V, Louis J, Ayre BG, Reese JC, Pegadaraju V, Shah J. 2011. TREHALOSE PHOSPHATE SYNTHASE11-dependent trehalose metabolism promotes Arabidopsis thaliana defense against the phloem-feeding insect Myzus persicae. Plant J 67(1): 94–104.

Song J, Win J, Tian M, Schornack S, Kaschani F, Ilyas M, van der Hoorn RA, Kamoun S. 2009. Apoplastic effectors secreted by two unrelated eukaryotic plant pathogens target the tomato defense protease Rcr3. Proc Natl Acad Sci U S A 106(5): 1654–1659.

Stam R, Howden AJ, Delgado-Cerezo M, M M Amaro TM, Motion GB, Pham J, Huitema E. 2013a. Characterization of cell death inducing Phytophthora capsici CRN effectors suggests diverse activities in the host nucleus. Front Plant Sci 4: 387.

Stam R, Jupe J, Howden AJ, Morris JA, Boevink PC, Hedley PE, Huitema E. 2013b. Identification and Characterisation CRN Effectors in Phytophthora capsici Shows Modularity and Functional Diversity. PLoS One 8(3): e59517.

Stam R, Motion GB, Boevink P, Huitema E. 2013c. A conserved oomycete CRN effector targets and modulates tomato TCP14-2 to enhance virulence. bioRxiv.

Thorpe P, Cock PJA, Bos J. 2016. Comparative transcriptomics and proteomics of three different aphid species identifies core and diverse effector sets. Bmc Genomics 17.

Thorpe P, Escudero-Martinez CM, Cock PJA, Eves-van den Akker S, Bos JIB. 2018. Shared transcriptional control and disparate gain and loss of aphid parasitism genes. Genome Biology and Evolution.

van den Burg HA, Kini RK, Schuurink RC, Takken FL. 2010. Arabidopsis small ubiquitin-like modifier paralogs have distinct functions in development and defense. Plant Cell 22(6): 1998–2016.

Verma V, Croley F, Sadanandom A. 2018. Fifty shades of SUMO: its role in immunity and at the fulcrum of the growth-defence balance. Mol Plant Pathol 19(6): 1537–1544.

Wang N, Zhao P, Ma Y, Yao X, Sun Y, Huang X, Jin J, Zhang Y, Zhu C, Fang R, et al. 2019. A whitefly effector Bsp9 targets host immunity regulator WRKY33 to promote performance. Philos Trans R Soc Lond B Biol Sci 374(1767): 20180313.

Wang Y, Bouwmeester K, van de Mortel JE, Shan W, Govers F. 2013. A novel Arabidopsis-oomycete pathosystem: differential interactions with Phytophthora capsici reveal a role for camalexin, indole glucosinolates and salicylic acid in defence. Plant Cell Environ 36(6): 1192–1203.

Weßling R, Epple P, Altmann S, He Y, Yang L, Henz SR, McDonald N, Wiley K, Bader KC, Gläßer C, et al. 2014. Convergent targeting of a common host protein-network by pathogen effectors from three kingdoms of life. Cell Host Microbe 16(3): 364–375.

Xu HX, Qian LX, Wang XW, Shao RX, Hong Y, Liu SS. 2019. A salivary effector enables whitefly to feed on host plants by eliciting salicylic acid-signaling pathway. Proc Natl Acad Sci U S A 116(2): 490–495.

Yachdav G, Kloppmann E, Kajan L, Hecht M, Goldberg T, Hamp T, Hönigschmid P, Schafferhans A, Roos M, Bernhofer M, et al. 2014. PredictProtein--an open resource for online prediction of protein structural and functional features. Nucleic Acids Res 42(Web Server issue): W337–343.

Yang S, Hua J. 2004. A haplotype-specific Resistance gene regulated by BONZAI1 mediates temperature-dependent growth control in Arabidopsis. Plant Cell 16(4): 1060–1071.

Zhao Q, Xie Y, Zheng Y, Jiang S, Liu W, Mu W, Liu Z, Zhao Y, Xue Y, Ren J. 2014. GPS-SUMO: a tool for the prediction of sumoylation sites and SUMO-interaction motifs. Nucleic Acids Res 42(Web Server issue): W325–330.

